# Tau local structure shields amyloid motif and controls aggregation propensity

**DOI:** 10.1101/330266

**Authors:** Dailu Chen, Kenneth W. Drombosky, Zhiqiang Hou, Levent Sari, Omar M. Kashmer, Bryan D. Ryder, Valerie A. Perez, DaNae R. Woodard, Milo M. Lin, Marc I. Diamond, Lukasz A. Joachimiak

**Author notes:** These authors contributed equally (Dailu Chen, Kenneth W. Drombosky). current address: Mylan NV, 3711 Collins Ferry, Morgantown, WV 26505.

## Abstract

Tauopathies are neurodegenerative diseases characterized by intracellular amyloid deposits of tau protein. Missense mutations in the tau gene (*MAPT*) correlate with aggregation propensity and cause dominantly inherited tauopathies, but their biophysical mechanism driving amyloid formation is poorly understood. Many disease-associated mutations localize within tau’s repeat domain at inter-repeat interfaces proximal to amyloidogenic sequences, such as ^306^VQIVYK^311^. Using cross-linking mass spectrometry, intramolecular FRET, recombinant protein and synthetic peptide systems, *in silico* modeling, and cell models, we conclude that the aggregation prone ^306^VQIVYK^311^ motif forms metastable compact structures with the upstream sequence that modulates aggregation propensity. Disease-associated mutations, isomerization of a critical proline, or alternative splicing are all sufficient to destabilize this local structure and trigger spontaneous aggregation. These findings provide a biophysical framework to explain the basis of early conformational changes that may underlie genetic and sporadic tau pathogenesis.

## INTRODUCTION

Tauopathies comprise a group of over 20 neurodegenerative diseases in which tau protein aggregates in neurons and glia. Tau aggregation correlates strongly with the degree of dementia and neurodegeneration, especially in Alzheimer’s Disease. The mechanisms by which disease-associated mutations, alternative splicing, or other events promote aggregation and pathology is not well understood. Understanding the molecular basis of tau aggregation could greatly improve diagnosis and treatment of tauopathies.

The N-terminal ∼200 and C-terminal ∼80 residues of tau are largely disordered, rendering this system refractory to high-resolution studies using structural biology methods^1^. In contrast, the tau repeat domain (tau RD), which spans residues 243 to 365, is predicted to be more structured^2^, forms the core of amyloid fibrils^3^, and is the minimal region to propagate tau prion strains^4^. Tau RD contains an amyloid motif (^306^VQIVYK^311^) (**Figure 1 A**) that is central to conversion between the soluble and insoluble states, as it mediates self-assembly, drives amyloid formation *in vitro*^5^ and promotes pathology *in vivo*^6^. Nuclear Magnetic Resonance (NMR) experiments on tau indicate that in solution the ^306^VQIVYK^311^ motif adopts a β-strand conformation^2,7^. Recent cryo-EM studies of tau patient-derived fibrils have shown that ^306^VQIVYK^311^ mediates important contacts in these structures^3,8^. Despite these structural studies, it’s not clear how native tau avoids aggregation, nor is it clear how tau transitions from a soluble state to an aggregated assembly.

**Figure 1.**
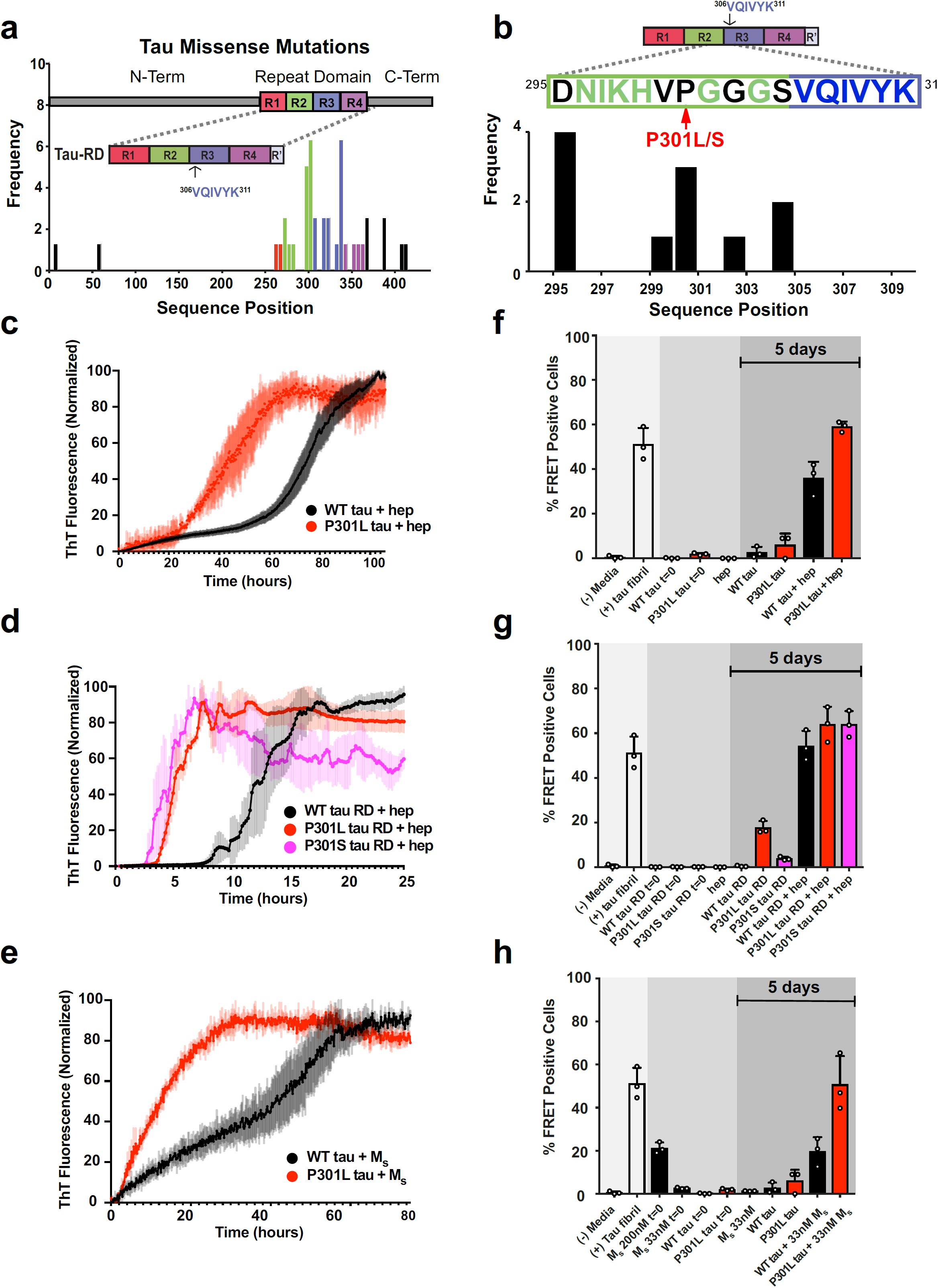
Tauopathy mutations cluster to inter-repeat regions. **a.** Disease-associated mutation frequency found in human tauopathies. Most mutations are found within the repeat domain (tau RD) (repeat 1 = red; repeat 2 = green; repeat 3 = blue; repeat 4 = purple). Amyloidogenic sequence ^306^VQIVYK^311^ is shown in the inset cartoon. **b.** Detailed mutation frequencies found near the ^306^VQIVYK^311^ amyloid motif. **c.** FL WT tau and mutant P301L tau at a 4.4 µM concentration were mixed with stoichiometric amounts of heparin (4.4 µM), and allowed to aggregate in the presence of ThT at room temperature. Control WT and P301L tau in the absence of heparin yielded no detectible ThT signal change (less than 2-fold ratio to background signal) over the course of the experiment (see Supplemental Table 1). ThT fluorescence was normalized to the maximum for each condition. **d**. WT tau RD and mutant P301L and P301S tau RD at a 4.4 µM concentration were each mixed with equimolar amounts of heparin (4.4 µM), and allowed to aggregate in the presence of ThT at room temperature. Control WT, P301L and P301S tau RD in the absence of heparin yielded no detectible ThT signal change (less than 2-fold ratio to background signal) over the course of the experiment (see Supplemental Table 1). **e**. WT FL tau and mutant P301L tau at a 4.4 µM concentration were mixed with substoichiometric M_s_ tau seed (33nM) and allowed to aggregate in the presence of ThT at room temperature. Control WT and P301L tau in the absence of M_s_ yielded no detectible ThT signal change (less than 2-fold ratio to background signal) over the course of the experiment (see Supplemental Table 1). All ThT experiments were carried out in triplicate. The data are shown as the average with standard deviation and are colored according to mutation. **f-h**. After 120 hours of *in vitro* incubation, proteins from previous ThT experiments were transduced into tau biosensor cells *via* lipofectamine (**Methods**). FRET signal from each condition (tau RD-CFP/tau RD-YFP) was measured by flow cytometry on 3 biological triplicates of at least 10,000 cells per condition. Error bars represent a 95% CI of each condition.

Polyanions such as heparin, nucleic acids, and arachidonic acid are commonly used to induce tau aggregation *in vitro*^9-11^. Solution NMR experiments mapped the tau-heparin binding site to repeat 2 just prior to the ^306^VQIVYK^311^ motif, but how this binding event modulates tau aggregation remains unclear^12^. Double electron-electron resonance (DEER) experiments indicated an expansion of this region upon heparin binding^9^. Cryo-EM structures also suggested an extended conformation of tau when bound to tubulin^13^. Other work mapping the recruitment of molecular chaperones to tau indicated that many chaperones, including Hsp40, Hsp70 and Hsp90, localize around ^306^VQIVYK^311 14^. Furthermore, unfolding of tau RD appeared to promote chaperone binding to the amyloid motif, suggesting that local conformational changes may help recruit factors to limit aggregation^15^. Recent data from our group indicated that soluble monomeric tau exists in at least two conformational ensembles: inert monomer (M_i_) which does not spontaneously self-assemble, and seed-competent monomer (M_s_) which spontaneously self-assembles into amyloid^16^. M_s_ itself adopts multiple stable structures that encode different tau prion strains^17^, which are unique amyloid assemblies that faithfully replicate in living systems. Based on extrapolations, the existence of an aggregation prone monomer of tau had been previously proposed ^18,19^ but our study was the first to biochemically isolate and characterize this species^16^. Different forms of M_s_ have been purified from recombinant protein, and tauopathy brain lysates^16,17^. Using multiple low-resolution structural methods, we have mapped critical structural changes that differentiate M_i_ from M_s_ to near the ^306^VQIVYK^311^ motif and indicated that the repeat 2 and 3 region in tau is extended in M_s_, which exposes the ^306^VQIVYK^311^ motif^16^. In contrast, intramolecular disulfide bridge between two native cysteines that flank ^306^VQIVYK^311^ in tau RD is predicted to form a local structure that is incompatible with the formation of amyloid^20^. Thus, conformational changes surrounding the ^306^VQIVYK^311^ amyloid motif appear critical to modulate aggregation propensity. A fragment of tau RD in complex with microtubules hinted that ^306^VQIVYK^311^ forms local contacts with upstream flanking sequence^21^. This was recently supported by predicted models guided by experimental restraints from crosslinking mass spectrometry^16^ and is consistent with independent NMR data^22,23^.

Based on our prior work^16^ we hypothesized that tau adopts a β-hairpin that shields the ^306^VQIVYK^311^ motif and that disease-associated mutations near the motif may contribute to tau’s molecular rearrangement which transforms it from an inert to an early seed-competent form by perturbing this structure. Many of the missense mutations genetically linked to tau pathology in humans occur within tau RD and cluster near ^306^VQIVYK^311 24^ (**Figure 1 a, b**), such as P301L and P301S. These mutations have no definitive biophysical mechanism of action, but are nevertheless widely used in cell and animal models^25,26^. Solution NMR experiments on tau RD encoding a P301L mutation have shown local chemical shift perturbations surrounding the mutation resulting in an increased β-strand propensity^27^. NMR measurements have yielded important insights but require the acquisition of spectra in non-physiological conditions, where aggregation is prohibited. Under these conditions weakly populated states that drive prion aggregation and early seed formation may not be observed^28^.

As with disease-associated mutations, alternative splicing also changes the sequence N-terminal to ^306^VQIVYK^311^. Tau is expressed in the adult brain primarily as two major splice isoforms: 3-repeat and 4-repeat^29^. The truncated 3-repeat isoform lacks the second of four imperfectly repeated segments in tau RD. Expression of the 4-repeat isoform correlates with the deposition of aggregated tau tangles in many tauopathies^30^ and non-coding mutations that increase preferential splicing or expression of the 4-repeat isoform cause dominantly inherited tauopathies^30-32^. It is not obvious why the incorporation or absence of the second repeat correlates with disease, as the primary sequences, although imperfectly repeated, are relatively conserved.

Previous reports have focused on the structure of a repeat with the assumption that each repeat functions independently within tau RD^33^. These have described a relationship between the length of a repeat fragment, its propensity to spontaneously aggregate, and its seeding capacity in cells^33^. However, inter-repeat interactions may also influence aggregation given that both alternative splicing and many disease-associated mutations cluster around the repeat interfaces (**Figure 1 a**). Our prior work suggested that wild-type tau aggregates less efficiently because the flanking sequence shields ^306^VQIVYK^311 16^. We hypothesize that the intrinsically disordered tau protein evolved to minimize aggregation by adopting local structure that shields the ^306^VQIVYK^311^ amyloid motif from interactions leading to seed formation and amyloid propagation. We predict that disease-associated mutations, alternative splicing, or other factors can destabilize this local structure and expose ^306^VQIVYK^311^ leading to self-assembly. Therefore, we employed an array of *in silico, in vitro*, and cellular assays to elucidate the molecular interactions and physiological consequences of ^306^VQIVYK^311^ within tau.

## RESULTS

### Disease-associated mutations at P301 promote rapid aggregation *in vitro* and in cells

Missense mutations that change proline 301 to leucine or serine cause dominantly inherited tauopathy^34^ and are associated with neurodegeneration in model systems^26,35^, though the biophysical mechanism is not understood. We studied changes in aggregation propensity driven by mutations at P301 in full-length (FL) tau (2N4R; amino acids 1-441) and tau repeat domain (tau RD; amino acids 244-380). First, we monitored aggregation of FL wild-type (WT) tau and mutant (P301L) tau using a Thioflavin T (ThT) fluorescence assay induced with stoichiometric amounts of heparin. We observed that P301L tau (t_1/2_ = 41.6 ± 0.5 hrs) aggregated more rapidly compared to WT tau (t_1/2_ = 75 ± 0.3 hrs) (**Figure 1 c**). Next, we compared changes in heparin-induced aggregation of the tau RD, comparing WT, P301L and P301S mutants. We again observed that the two mutants aggregated faster (P301L tau RD, t_1/2_ = 5.2 ± 0.1 hrs; P301S tau RD, t_1/2_ = 3.9 ± 0.1 hrs) than WT tau RD (WT tau RD, t_1/2_ = 12.5 ± 0.2 hrs) (**Figure 1 d**). Consistently, we found that mutations at position 301 (from proline to either leucine or serine) increased aggregation rates by ∼2-fold compared to WT in both FL tau and tau RD constructs. Thus, *in vitro,* tau RD recapitulates key aspects of aggregation observed in FL tau.

The inert conformation of monomeric tau (M_i_) requires cofactors, such as heparin, to spontaneously aggregate *in vitro*, while the seed-competent monomer (M_s_), derived from recombinant protein or Alzheimer’s patient brain material, readily self-assembles to form amyloid^16^. Previously we determined that M_s_ converts FL tau into fibrils at sub-stoichiometric ratios, in contrast to the stoichiometric amounts necessary in heparin-containing reactions^16^. In this study, we evaluated the aggregation propensity of the P301L mutant compared to WT when incubated in the presence of recombinantly produced M_s_. We incubated FL tau with sub-stoichiometric amounts of M_s_ (1:133) and monitored aggregation using ThT. In comparison, we observed that M_s_-seeded P301L tau self-assembled more rapidly (P301L tau, t_1/2_ = 8.5 ± 0.6 hrs) than the WT protein (WT tau, t_1/2_ = 40 ± 1.1 hrs) (**Figure 1 e).** P301L tau aggregated faster than WT tau with a 4-fold increase in rate after seeding by M_s_. Independent of induction – heparin or M_s_ – P301L assembled into ThT positive aggregates more rapidly. Moreover, tau appeared to be more sensitive to M_s_ seeded aggregation compared to heparin, given the sub-stoichiometric ratios needed for robust aggregation. Based on these observations we next sought to determine why M_s_ seeds appear to be more efficient than heparin at converting tau into fibrils, and why mutations at P301 increase tau’s sensitivity to seeded aggregation.

We evaluated the change in conformation between M_i_ and M_s_ using fluorescence resonance energy transfer (FRET). Tau encodes two cysteine residues at positions C291 in repeat 2 and C322 in repeat 3 (**Methods**, **Table 1, Supplemental Figure 1 a**), which serendipitously flank the amyloid motif ^306^VQIVYK^311^. We labeled M_i_ and M_s_ at C291 and C322 with donor and acceptor fluorophores using maleimide chemistry and measured their steady-state FRET intensities. M_i_, which is predicted to be in a more closed conformation^16^, showed higher FRET efficiencies (E = 0.58 ± 0.12) compared to M_s_ (E = 0.25 ± 0.078) (**Supplemental Figure 1 b**). Denaturing M_i_ and M_s_ at 95 °C overnight yielded similar FRET efficiencies (E = 0.24 ± 0.056, E = 0.29 ± 0.04, respectively) (**Supplemental Figure 1 b**). Importantly, FRET efficiencies calculated for free dyes were low and remained constant at low and high temperatures (**Supplemental Table 2**). FRET efficiencies for both 95 °C denatured proteins were only modestly above the background FRET efficiency for the free dye negative control (**Supplemental Table 2**), suggesting a minimal interaction between these two regions. Furthermore, FRET from M_s_ was reduced even at room temperature, likely reflecting a similarly extended conformation of the repeat 2 and 3 region versus a relatively compact conformation in M_i_. Thus, the effectiveness of M_s_ to seed aggregation of M_i_ may be explained by a direct change in conformation proximal to the ^306^VQIVYK^311^ motif, which is central to tau aggregation. This suggested that mutations at the P301 may exacerbate aggregation by unfolding the region surrounding the ^306^VQIVYK^311^ motif, thereby producing a more compatible conformation for the similarly expanded aggregation-prone M_s_ seed.

**Table 1:**
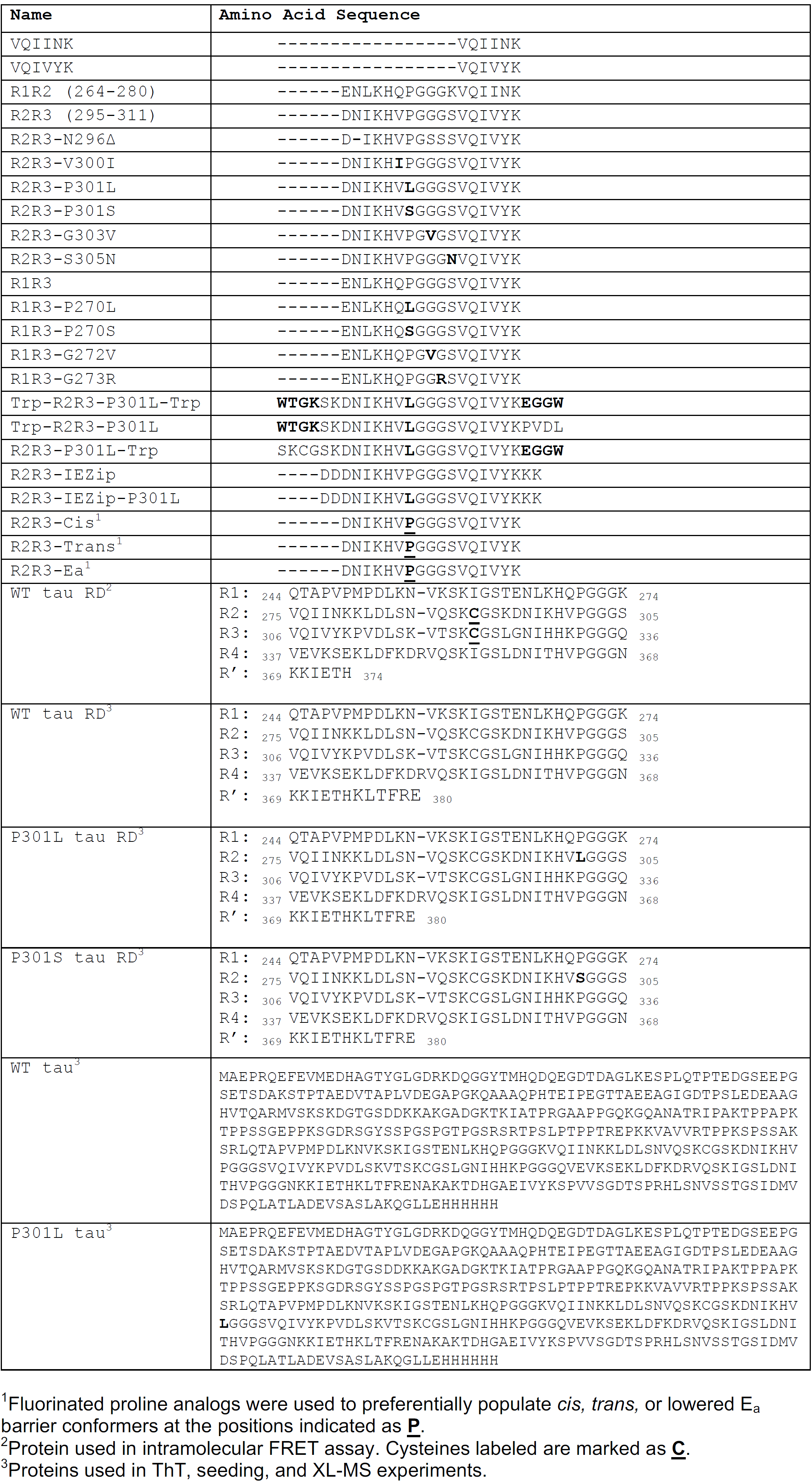
List of Peptide and Protein Sequences

**Table 2:**
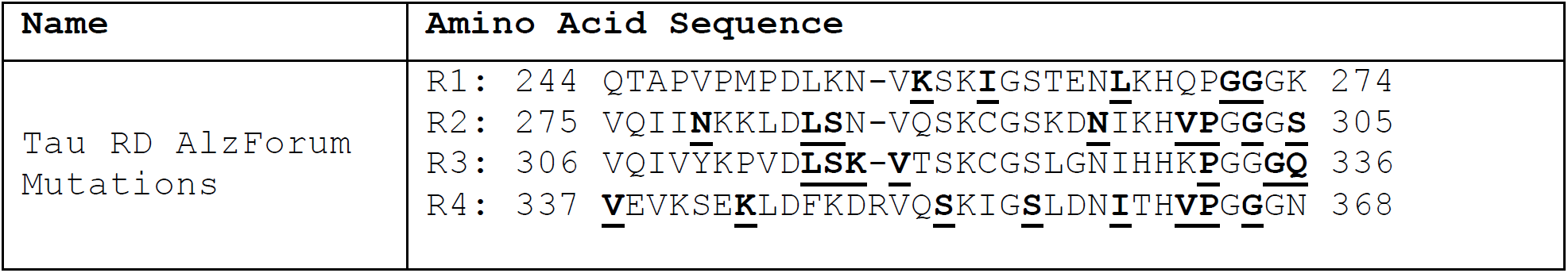
List of AlzForum Disease-Associated Mutations

To test the structural compatibility of aggregates formed by *in vitro* tau models, we employed tau biosensor HEK293 cells that stably express tau RD (P301S) fused to cyan or yellow fluorescent proteins ^25^. These cells sensitively report a FRET signal (tau RD-CFP/tau RD-YFP) only when aggregated in response to tau amyloid seeds, and are unresponsive to aggregates formed by other proteins, such as huntingtin or α-synuclein^36^. Each sample formed amyloid fibril morphologies confirmed by transmission electron microscopy, except for samples not incubated with heparin or M_s_ and the low-concentration M_s_, where no large ordered structures were found (**Supplemental Figure 2**). The tau biosensor cells responded to FL tau fibrils created by exposure to heparin and showed an increase in seeding activity for the P301L mutant compared to WT fibrils (**Figure 1 f**). Next, we compared seeding for the tau RD heparin-induced fibrils and again found that P301L and P301S mutants produced higher seeding activity relative to WT (**Figure 1 g**). Lastly, the seeding activity for the M_s_-induced FL tau fibrils showed a 2-fold higher activity for P301L compared to WT (**Figure 1 h**). WT FL tau and tau RD control samples (no heparin or M_s_) did not produce seeding activity in cells while P301 mutants, both FL and tau RD, showed hints of seeding activity despite not yielding ThT signal *in vitro* (data not shown), perhaps due to the formation of oligomers not captured by ThT. As expected, 33nM M_s_ control exhibited seeding activity at the onset and did not change after 5 days, but overall signal was low due to the low concentrations used in the aggregation experiments. Interestingly, WT tau induced with 33nM M_s_ seeded at similar levels to concentrated control (200nM) M_s_ samples highlighting efficient conversion of WT tau into seed-competent forms (**Figure 1 h**). Thus, P301 mutations promote aggregation *in vitro* and in cells across different constructs. Importantly, these effects are conserved between FL tau and tau RD.

### Mutation at P301 destabilizes native tau structure

To determine how the P301L mutation drives conformational changes we labeled WT tau RD and P301L tau RD with fluorophores at C291 and C322, flanking the ^306^VQIVYK^311^ motif (**Figure 2 a**), and monitored FRET in a heat denaturation time-course experiment. FRET efficiency for WT tau RD gradually decreased at 65°C or 75°C, consistent with denaturation of a metastable local structure (**Figure 2 b**). We next evaluated P301L tau RD and found no detectible difference between monomeric WT tau RD FRET at physiological temperature (37°C). However, unlike WT, P301L tau RD rapidly lost FRET signal with heat denaturation (**Figure 2 c**), indicating perturbation of local structure. The magnitude of destabilization was consistent with the differences observed for seeded elongation (**Figure 1 g**) which may not be readily detected by global ensemble measurements like NMR.

**Figure 2.**
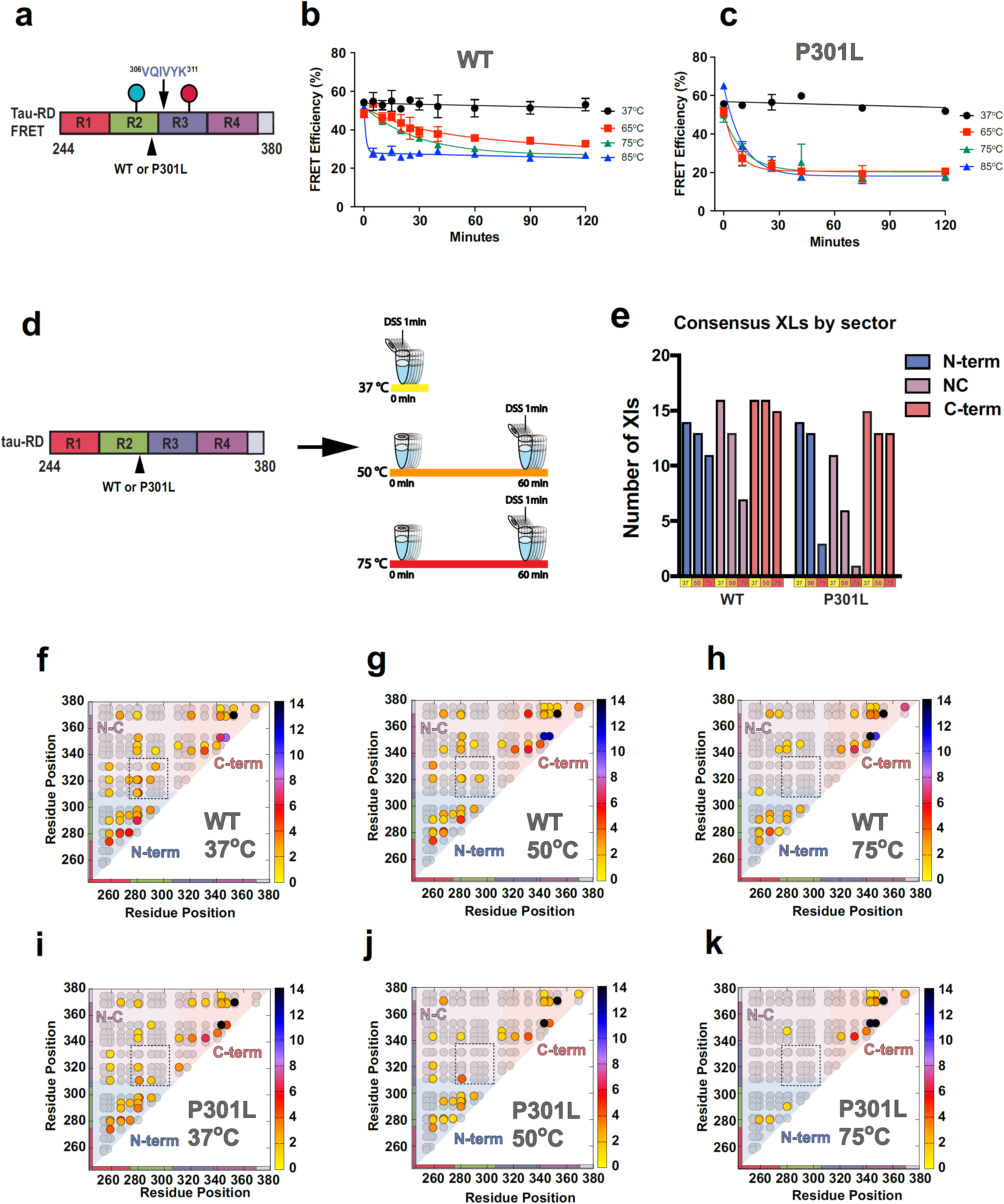
Tau RD encodes global and local structure. **a**. Cartoon schematic of tau RD used for FRET studies. For FRET studies, endogenous cysteines are labelled with Alexa-488 and Alexa-647 (red and blue circles), which flank the ^306^VQIVYK^311^ amyloid motif. **b-c**. FRET was measured for dye modified monomeric WT and P301L tau RD at a series of temperatures over a time course. **d**. Cartoon schematic of tau RD used for XL-MS studies colored according to repeat domain. Recombinant WT and P301L tau RD were heated at 37°C, 50°C or 75°C for one hour, then chemically crosslinked using DSS. After cross-linking, trypsin fragmentation, and LC-MS/MS analysis were performed. Each sample was carried out in 5 technical replicates. **e**. Total consensus crosslinks parsed by temperature and location in WT and P301L tau RD: within N-terminus (blue; residues 243-310; N-term), within C-terminus (orange; residues 311-380; C-term) and span N- and C-terminus (magenta; between residues 243-310 and 311-380; N-C). **f-h**. Consensus crosslinks (circles) are shown in contact maps color coded by average frequency across replicates. The theoretical lysine pairs are shown in the background as gray circles. Crosslink contacts within the N-term (blue), C-term (red) and across N-to C-term (purple) are shown as sectors. The x- and y-axis are colored according to repeat number as in Figure 1. The dashed boxes define inter-repeat crosslinks observed between repeat 2 and repeat 3. **i-k**. Same as f-h above, except with tau RD that contains a P301L mutation.

We next used an orthogonal approach to test these ideas. Cross-linking mass spectrometry (XL-MS) defines amino acid contacts in proteins and can thus guide the determination of structure for large complexes, transient interactions and dynamics of intrinsically disordered proteins ^16,37,38^. To preclude the formation of intermolecular crosslinks between monomers, low concentration samples of WT, P301L and P301S tau RD were incubated at different temperatures, reacted with disuccinimidyl suberate (DSS; a primary amine crosslinker) for 1 minute and quenched (**Figure 2 d**). The crosslinked protein monomers were confirmed by SDS-PAGE (**Supplemental Figure 3 a**). Cross-linked samples were trypsin digested, analyzed by mass spectrometry and the spectra were searched using the Xquest^39^ to identify intramolecular protein contact pairs (**Methods and Supplemental Table 2**). In each dataset, the cross-links reported represent consensus data across five independent samples with a low false discovery rate (**Methods, Supplemental Table 2 and 3 and Supplemental Figure 4**). XL-MS of recombinant WT tau RD acquired at 37°C revealed three clusters of interactions that localize within the N-terminus (residues 243-310; N-term), within the C-terminus (residues 311-380; C-term) and span N- and C-termini (N-C; between residues 243-310 and 311-380) (**Figure 2 e, f**). Importantly, the experimentally observed crosslinks represent only a small subset of all theoretical lysine pairs suggesting that tau RD samples discrete modes of contacts (**Figure 2 f, grey circles**). Heating the samples to 50°C or even 75°C decreased the number of N-C long-range and N-term short-range contacts identified (**Figure 2 e, g, h and Supplemental Figure 3 b).** The data acquired for WT tau RD in physiological conditions is consistent with a loose metastable structure comprised of weak long-range and short-range contacts that are sensitive to temperature.

In contrast, XL-MS of recombinant tau RD with the P301L mutation revealed an increased susceptibility to heat denaturation. At 37°C, the crosslinks found in P301L tau RD (**Figure 2 i**) were similar in pattern to WT, except for fewer N-C terminal long-range contacts **(Figure 2 e and Supplemental Figure 3 b, c)**. However, samples incubated at 50°C and 75°C revealed a significant reduction in both long-range and short-range contacts in P301L tau RD compared to WT **(Figure 2 e)**. The loss of short-range contacts, both the total number of crosslinks and the abundance of each crosslink, were detected particularly within the N-terminal sector, which sits upstream of P301 (**Figure 2 e, i-k and Supplemental Figure 3 b, c**). In contrast, the C-term local contacts sample many theoretical lysine-lysine pairs and remained relatively constant across temperatures for both WT and P301L tau RD possibly suggesting a higher degree of disorder that is independent of the mutation site. Moreover, a stepwise loss of the inter-repeat crosslinks between repeat 2 and 3 was seen in the heat denaturation of WT tau RD and was more pronounced with the P301L mutation, indicating an unfolding of local structure at the interface of repeat 2 and 3, encompassing ^306^VQIVYK^311^ (**Figure 2 f-k, inset box**). Thus, while the number of crosslinks identified was comparable between WT and P301L tau RD at 37°C, P301L tau RD retained approximately half as many crosslinks as WT when heated. Similar crosslinking profiles were observed for the P301S tau RD sample (**Supplemental Figure 3 d**). Thus, the XL-MS data were consistent with findings from FRET labeling: P301L tau RD lacks the thermostability observed in WT tau RD.

### *In silico* modeling of tau RD shows inter-repeat elements adopt local structure

To gain insight into what types of local structures the inter-repeat elements can form, we used ROSETTA to predict structures of tau RD. We built 5,000 models using two parallel strategies in ROSETTA: *ab initio* which employed fragment libraries derived from experimental structures^40^ and CS-ROSETTA which leveraged available chemical shifts for tau RD to produce fragment libraries^41^. Both approaches led to a diversity of models consistent with experimentally determined radii of gyration^42^ **(Supplemental Figure 5 a-d)**. Analysis of each structural ensemble showed a propensity to form hairpin-like structures across R1R2, R2R3, R3R4 and R4R’ repeat interfaces centered on the ^271^PGGG^274^, ^301^PGGG^304^, ^333^PGGG^336^ and ^366^PGGG^369^ sequences (**Supplemental Figure 5 e and 6**). Previously published solution NMR data have shown that the PGGG sequences in tau can adopt type II β-turns^7^, and the ^301^PGGG^304^ sequence preceding ^306^VQIVYK^311^ is compatible with the formation of a β-hairpin. We illustrated the R2R3 ^306^VQIVYK^311^-containing fragment derived from low energy expanded models produced by each method (**Supplemental Figure 5 c, d**). The ^306^VQIVYK^311^-containing interface has the highest frequency of disease-associated mutations, particularly P301L and P301S (**Figure 1 a**). Other potential amyloid-forming regions, such as ^275^VQIINK^280^, which is capable of aggregation (**Supplemental Figure 7**), is also preceded by ^271^PGGG^274^ and predicted to form a β-hairpin (**Supplemental Figure 5 d and Supplemental Figure 6**), however, it is absent in recent cryo-EM structures of tau aggregates^3,8^. Mapping known missense mutations onto the *ab initio* β-hairpin structure at R2R3 interface (**Supplemental Figure 5 f**), we hypothesized that this cluster of disease-associated mutations could destabilize the β-hairpin secondary structure, thus exposing the amyloid motif ^306^VQIVYK^311^ and enabling aggregation. This model is compatible with recent cryo-EM findings that indicate a disengagement of the ^306^VQIVYK^311^ N-terminal flanking sequence in a fibril structure^3^. Thus, we focused our studies on the R2R3 motif of tau that contains ^306^VQIVYK^311^.

### P301L promotes extended forms of tau

*In silico* modelling corroborated recent biochemical findings^16^ and suggested a minimal sequence necessary to form a collapsed structure around ^306^VQIVYK^311^. To understand how these structures might self-assemble, we employed molecular dynamics (MD) simulations of two tau peptide fragments comprising the minimally structured fragment centered around the R2R3 interface (^295^DNIKHVPGGGSVQIVYK^311^): R2R3-WT and R2R3-P301L (**Table 1 and Figure 4 b**). To enable sufficient sampling of oligomer structures, we employed an unbiased algorithm based on a recently-developed symmetry-constraint approach^43^. The trimer conformations obtained in simulations are depicted on a root-mean-square deviation (RMSD) matrix for both R2R3-WT (**Figure 3 a**) and the R2R3-P301L mutant peptide fragments (**Figure 3 b**). For R2R3-WT peptide fragment, we observed a dominant population of trimeric conformations composed of hairpins, while the P301L disease-associated mutation stabilized an extended fibrillar form. The energy basin for R2R3-WT peptide fragment was predicted to be 5 to 6 kJ/mol lower in a collapsed state than an extended state, whereas the R2R3-P301L peptide fragment was 3 kJ/mol lower in an extended state than a collapsed state (**Figure 3 c**). Additionally, the free energy surface suggested an energy barrier of approximately 5 kJ/mol to convert R2R3-WT peptide fragment from collapsed to extended. That same barrier however was less than 1 kJ/mole for the R2R3-P301L peptide fragment, suggesting a faster rate of kinetic conversion from collapsed hairpin to extended fibrillar form. Thus, MD predicts that P301L mutation promotes amyloid assembly by destabilizing monomeric hairpin structures.

**Figure 3.**
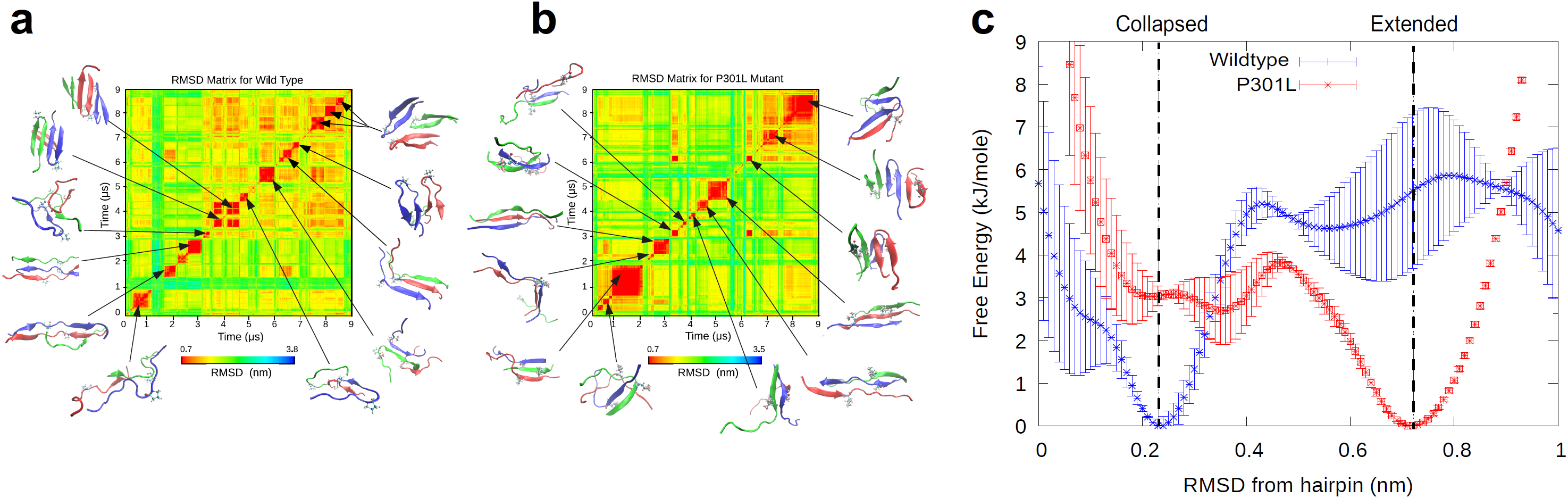
Wildtype and mutant peptides differentially populate collapsed and extended conformations. **a**. Conformations obtained for a trimer of WT peptide fragment (R2R3-WT) with the sequence ^295^DNIKHVPGGGSVQIVYK^311^. Two-dimensional root-mean-squared-differences (RMSD’s) are calculated between all pairs of conformations visited during MD simulations. Snapshots of trimeric structures are depicted for selected metastable basins, with each peptide monomer represented by a different color. **b**. The same analysis as above, but for the P301L substituted trimer. **c**. The free energy surface as a function of deviation from a canonical hairpin structure. Two distinct basins, corresponding to collapsed and extended sub-ensembles, are found in WT and P301L peptide fragment respectively.

### Tau amyloid formation is governed by flanking residues

In tau RD, ^306^VQIVYK^311^ is necessary for amyloid formation^5,6^. In solution, ^306^VQIVYK^311^ hexapeptide aggregates spontaneously and rapidly as measured by ThT fluorescence intensity (**Supplemental Figure 7**) whereas the upstream N-terminal sequence ^295^DNIKHV^300^ does not aggregate. To experimentally test the prediction of a local hairpin structure encompassing ^306^VQIVYK^311^, we employed a mutagenesis study on synthetic peptide systems that recapitulate the minimal hairpin sequence (**Table 1**). Consistent with the prediction from MD simulation, R2R3-WT peptide fragment did not aggregate readily, with no ThT detected within 96 hrs (**Figure 4 a, c**). By contrast, single disease-associated mutations (**Figure 4 b**) substituted into the R2R3 peptide fragment were sufficient to promote spontaneous amyloid formation: R2R3-P301S (t_1/2_ = 4.1 ± 1.3 hours), R2R3-P301L (t_1/2_ = 7.2 ± 0.2 hours), R2R3-N296Δ (t_1/2_ = 31.9 ± 0.2 hours), R2R3-G303V (t_1/2_ = 32.1 ± 0.7 hours), R2R3-S305N (t_1/2_ = 41.2 ± 0.2 hours), and R2R3-V300I (t_1/2_ = 77.8 ± 1.3 hours**, Figure 4 c**). Each of these peptides was confirmed to form amyloid-like fibril morphologies by transmission electron microscopy, except for the WT R2R3 peptide fragment where no large structures were found (**Figure 5 b-h**).

**Figure 4.**
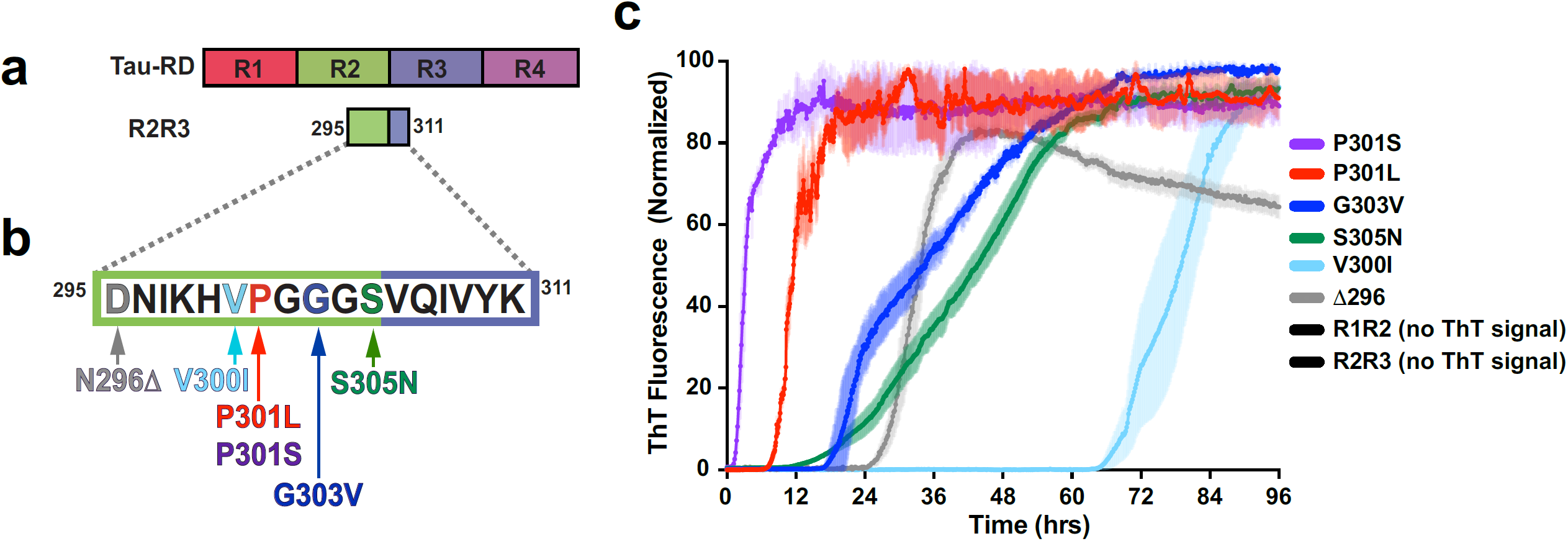
Tauopathy mutations drive aggregation propensity. **a**. Schematic of tau RD and the derived peptides representing the minimal structural element around ^306^VQIVYK^311^. **b**. WT and mutant peptides were disaggregated, resuspended to 200 µM, and allowed to aggregate in the presence of ThT at room temperature. WT R2R3 and R1R2 fragment peptides yielded no detectible ThT signal change (less than 2-fold ratio to background signal) over the course of the experiment (see Supplemental Table 1). ThT signals are shown as average of triplicates with standard deviation, are colored according to mutation and are normalized to the maximum for each condition.

**Figure 5.**
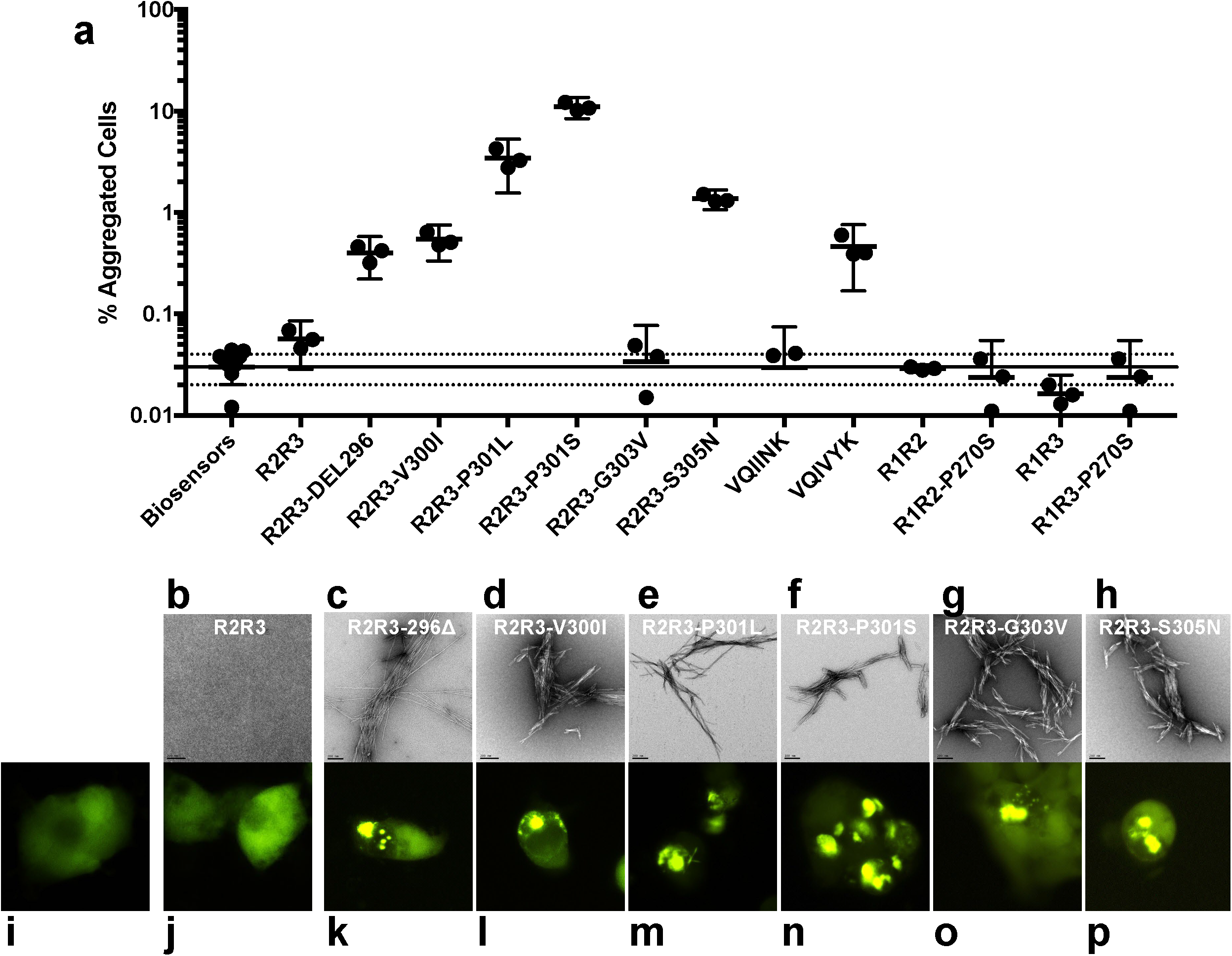
Peptides form amyloid structures and seed *in vivo*. **a**. After 96 hours of *in vitro* incubation, peptides from previous ThT experiments (**Figure 4 c**) were transduced into tau biosensor cells *via* lipofectamine (**Methods**). FRET signal from each condition (tau RD-CFP/tau RD-YFP) was measured by flow cytometry on 3 biological triplicates of at least 10,000 cells per condition. Error bars represent a 95% CI of each condition. Solid and dashed horizontal lines represent the mean and 95% error from untreated biosensor cells, respectively, for ease of statistical comparison. **b-h**. Electron microscopy images of each peptide from previous ThT experiments (**Figure 4 c**). The black bar represents 200 nm distance in each image **i-p**. Qualitative fluorescence microscopy images of tau biosensor cells immediately prior to Flow Cytometry experiments. i shows a representative image of untreated biosensor cells. j – p each shows a representative image of biosensor cells treated with samples from b-h respectively.

To test the structural compatibility of peptide aggregates formed by *in vitro* tau models, we again employed tau biosensor cells^25^. The tau biosensor cells responded to all disease-associated tau peptide fragments that aggregated spontaneously *in vitro*, but not to the wild-type R2R3 peptide fragment (which did not aggregate *in vitro*) (**Figure 5 a**). Qualitatively, biosensor cells retained their diffused tau localization when untreated or exposed to a wild-type R2R3 peptide fragment but formed fluorescent puncta when cultured with aggregated mutant peptides (**Figure 5 i-p**). Interestingly, the biosensor cells responded to disease-associated mutant peptides with varying degrees of sensitivity and created distinct aggregate morphologies. This is consistent with amyloid structures that act as distinct templates and form the basis of tau prion strains^4,44^. Thus, R2R3 peptide fragment model system responds to mutations *in vitro* and in cells similarly to the FL tau and tau RD system, suggesting that local conformational changes in tau can be recapitulated using shorter fragments.

### Tau splice variants reveal differential aggregation propensity

Tau is expressed in the adult brain as 6 major splice isoform types that include either 3 or 4 repeated segments within RD (**Figure 6 a**). 3R tau lacks the second of four imperfect repeats. 4R tau correlates strongly with aggregation in most tauopathies^30^ and mutations that increase splicing of the 4R isoform cause dominantly inherited tauopathies^30-32^. We examined whether this splice isoform affects the propensity of ^306^VQIVYK^311^-mediated aggregation due to the different composition of upstream flanking sequence. We constructed a series of peptide fragments to encompass the R1R3 interface (**Table 1**). This wild type peptide fragment R1R3 mimicking a 3R splice isoform did not spontaneously aggregate (data not shown). Surprisingly, an R1R3 peptide fragment with a corresponding P301L mutation (R1R3-P270L) also did not aggregate (**Figure 6**). We hypothesized that R1 leading sequence stabilizes the amyloid motif ^306^VQIVYK^311^ resulting in the aggregation resistance in the presence of disease-associated mutations.

**Figure 6.**
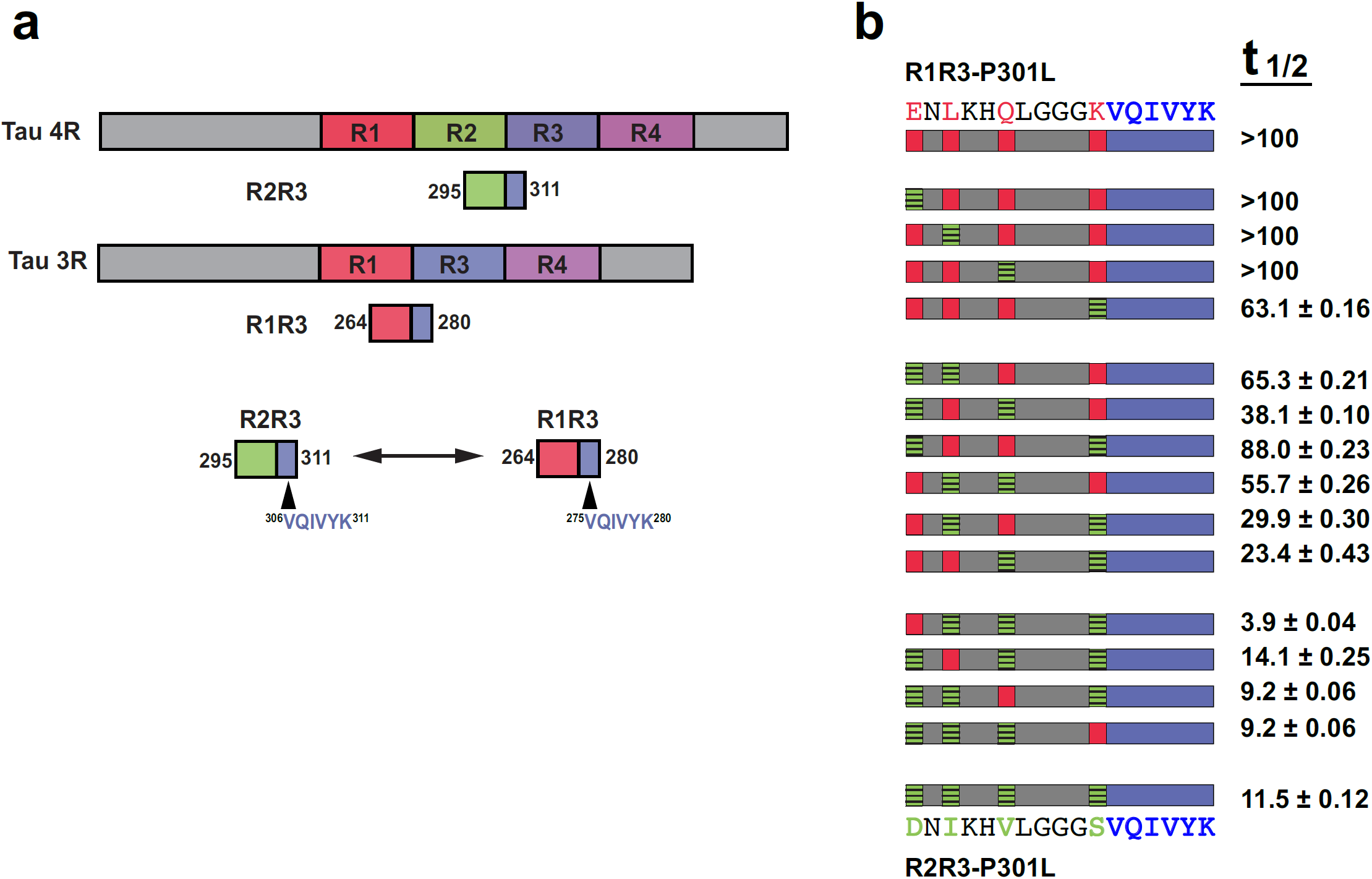
Alternative splicing modulates aggregation propensity. **a**. Cartoon schematic for tau 4R and 3R splice isoforms illustrate the difference in primary amino acid sequence leading into the amyloidogenic ^306^VQIVYK^311^ motif. **b**. A full combinatorial panel of R2R3-P301L and R1R3-P301L chimeras were aggregated *in vitro*. ^306^VQIVYK^311^ is shown in blue, amino acids common between the splice isoforms are shown in gray, amino acids unique to an R3 isoform are colored red, amino acids unique to an R4 isoform are colored green. The aggregation kinetics, represented as t_1/2_ in hours with 95% CI, are listed in the right-side column alongside its respective peptide.

The R1 leading sequence ^264^**<u>E</u>**N**<u>L</u>**KH**<u>Q</u>**PGGG**<u>K</u>**^273>^ differs from R2 ^295^**<u>D</u>**N**I**KH**<u>V</u>**PGGG**<u>S</u>**^304^ at four amino acid positions. To identify which amino acid(s) governed R1’s stronger inhibitory effects, we constructed 16 peptides with a P301L mutation to represent every combinatorial sequence between the two leading strands (**Figure 6 b**) and measured their aggregation kinetics (**Supplemental Figure 8)**. We identified a general trend where the R2R3-P301L peptide fragment aggregates in hours with zero or one R1 substitutions. With two R1 substitutions, R2R3-P301L peptide aggregation was delayed roughly an order of magnitude to tens of hours. With three R1 substitutions, R2R3-P301L peptide fragment aggregation was further delayed to hundreds of hours. With all four R1 substitutions in the peptide (R1R3-P301L), no ThT signal was observed within a week (**Figure 6 b, Supplemental Figure 8**). Thus, all four amino acids contributed to the ability of the R1 leading sequence to delay ^306^VQIVYK^311^-mediated spontaneous aggregation in a 3R splice isoform. This may explain the differential aggregation propensities of tau isoforms in human pathology.

### Stabilizing β-hairpin structure blocks P301L-mediated aggregation

Our model predicted shielding of the ^306^VQIVYK^311^ motif in tau *via* local β-structure, and thus we hypothesized that artificially stabilizing the termini of this local structure would promote a more inert, closed conformation. To test this, we flanked the R2R3-P301L peptide fragment with a tryptophan zipper (Trp-R2R3-P301L-Trp, **Table 1 and Figure 7 a**), which stabilizes a β-hairpin structure approximately −2.5 to −7 kJ/mol^45^. Consistent with our model, Trp-R2R3-P301L-Trp peptide fragment does not spontaneously aggregate *in vitro* (**Figure 7 b**). To ensure that this effect wasn’t a result of adding bulky tryptophan residues, we constructed control peptide fragments that contain only the N-term (Trp-R2R3-P301L) or the C-term (R2R3-P301L-Trp) portion of the tryptophan zipper sequence (**Figure 7 a**). Both half-sequence controls spontaneously aggregated, implying that a tryptophan at either position is insufficient to block aggregation (**Figure 7 b**). Only a fully intact tryptophan zipper that stabilizes a β-hairpin conformation ameliorates aggregation propensity. Alternative methods to stabilize a β-hairpin architecture, such as introducing isoelectric interactions, also delayed aggregation: peptides containing two additional aspartic acid on the N-terminus and two lysine on the C-terminus (R2R3-IEZip, **Table 1**) retarded R2R3-P301L peptide fragment aggregation over an order of magnitude (t_1/2_ = 7 hours to t_1/2_ > 70 hours, **Supplemental Figure 9**).

**Figure 7.**
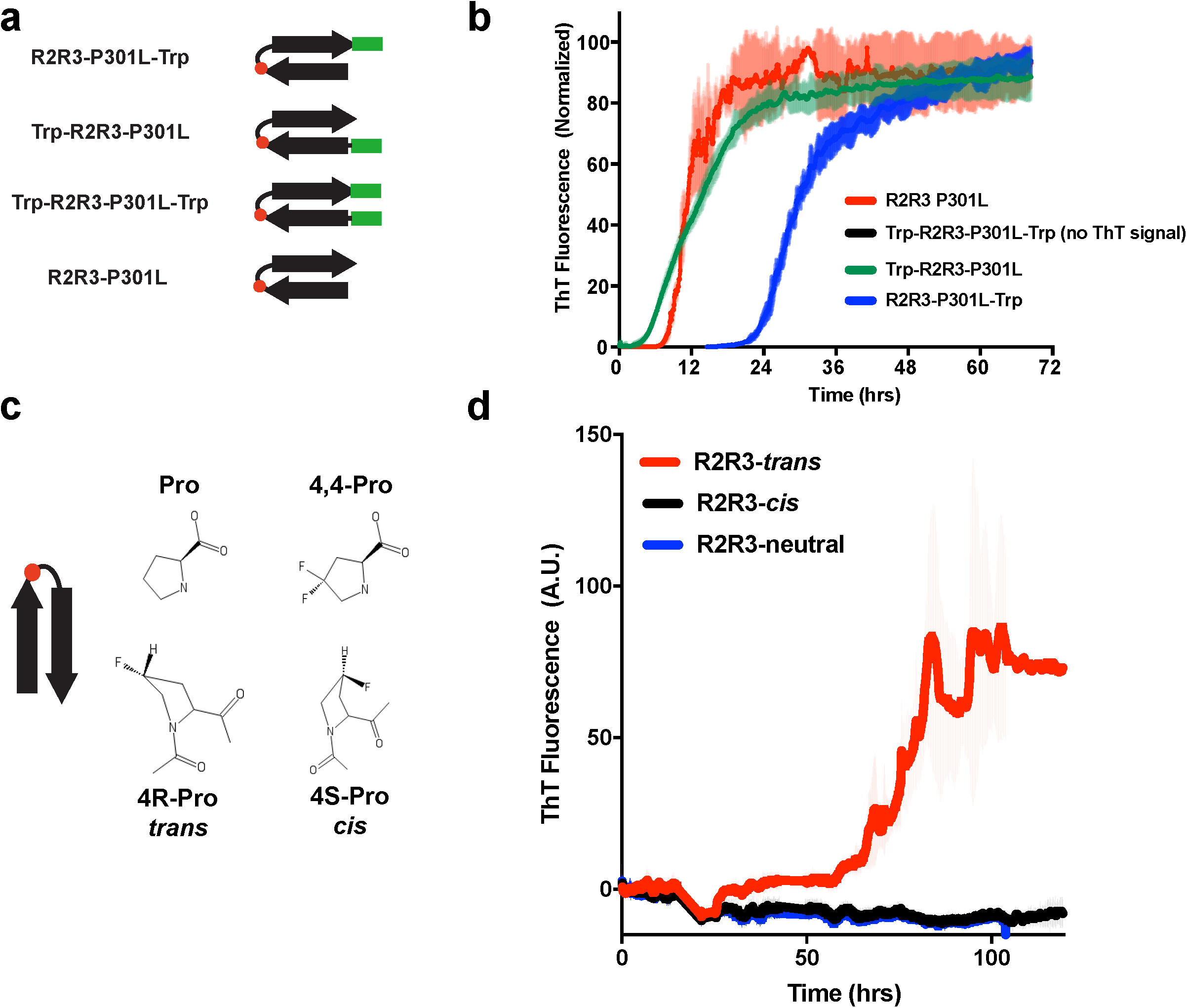
Enhancing β-hairpin structure rescues spontaneous aggregation phenotypes. **a**. Cartoon schematic representation of the tryptophan zipper motif (green bar) and controls used to stabilize a β-hairpin structure in an R2R3-P301L peptide fragment (**Table 1**). **b**. Aggregation reactions of the tryptophan zipper peptide and controls measured by ThT fluorescence. The Trp-R2R3-P301L-Trp fragment peptide yielded no detectible ThT signal change (less than 2-fold ratio to background signal) over the course of the experiment (see Supplemental Table 1) ThT signals are shown as average of triplicates with standard deviation and were normalized to the maximum for each condition. **c**. Schematic of proline and fluorinated proline analogs used to generate *cis* and *trans* proline conformers at the position corresponding to P301 (red circle) in peptide models. **d**. ThT aggregation reactions of the *cis, trans*, and neutral proline analogs substituted into the R2R3 peptide fragment. ThT signals are an average of 6 independent experiments with standard deviation shown.

To test this effect in cells, we engineered tau RD (P301S) biosensor cells encoding tryptophan zipper motifs that flank the R2R3 element. These biosensors had a significantly diminished capacity to be seeded; R2R3-P301S peptide fragment aggregates triggered aggregation in 14.5 ± 1.6% of tau biosensor cells, but only 0.3 ± 0.12% of the tryptophan zipper stabilized biosensor cells (**Supplemental Figure 10**).

### Proline 301 *cis-trans* isomerization modulates aggregation

Many proteins in the cell utilize proline isomerization as a molecular switch, such as heat shock protein activation^46^ or cell cycle regulation^47^. In some proteins, proline isomerization directly induces or mitigates aggregation into amyloid^48-50^. Proline isomerization events in tau have been proposed to play a role in aggregation and disease^48^, but P301 isomerization has not been linked to tau aggregation and pathology. With the fact that serine or leucine substitutions at P301 proximal to ^306^VQIVYK^311^ drastically alter aggregation propensity, we hypothesized that P301 plays a crucial role inducing a β-turn in a PGGG motif, which mediates a collapsed structure. To test whether isomerization of P301 could influence spontaneous amyloid formation, we constructed a series of R2R3 peptide fragments with proline analogs that preferentially populate either: (1) a *cis* rotamer (2S,4S)-fluoroproline; (2) a *trans* rotamer (2S,4R)-fluoroproline; or (3) an analog that easily interconverts between *cis* and *trans* (4,4)-difluoroproline (**Table 1, Figure 7 c**). Only R2R3-Trans peptide spontaneously aggregated (**Figure 7 d**), indicating the potential for proline isomerization events in tau pathogenesis.

## DISCUSSION

Here we establish the molecular and functional basis for how a series of prominent tau mutations drive aggregation. Integrating experimental and computational approaches, we independently and directly probed the local structural changes within tau. We identified metastable local structures within the inter-repeat junction of tau RD (the repeat 2-repeat 3 interface), which encompasses the amyloidogenic ^306^VQIVYK^311^ motif. This R2R3 interface becomes less stable when a disease-associated mutation is present, such as P301L, which is commonly employed in cell and animal models of tauopathy. Thus, P301L and similar mutations decrease the threshold for local structural expansion, especially in the presence of stressors (heat, seeds, heparin, or high concentration). This in turn is predicted to enhance the conversion of tau into a seed-competent form^16^. Thus, the proposed model rationalizes the fundamental molecular mechanisms of aggregation for P301L and at least 5 other mutations, explains why P301L spontaneously aggregates in animal and cellular models, and defines how splice isoforms of tau and proline isomerization at P301 may contribute to aggregation. Ultimately, these insights may inform the mechanisms of tauopathy in human disease and potential molecular targets for therapeutic development.

### Mechanism of tau seeding

*In vitro* induction of tau aggregation is typically achieved by the addition of polyanionic molecules such as heparin, arachidonic acid or nucleic acids^9-11^. It is thought that heparin binding to tau expands the local conformation of the repeat 2 and repeat 3 regions, thereby exposing amyloidogenic sequences for subsequent aggregation^9,12,16^. This process, however, requires stoichiometric amounts of polyanion and is not a physiological condition since heparin is not present intracellularly. Our recent work has elucidated a seed competent form of tau monomer that can promote tau aggregation. This seed competent monomeric tau is found in AD patient brains and is likely the incipient species contributing to pathology^16^. We find that sub-stoichiometric amounts of M_s_ (1:133) enhance the rate of WT tau aggregation relative to heparin. Parallel experiments with P301L tau show an even more dramatic enhancement. Our data support that the ^306^VQIVYK^311^ motif is preferentially exposed in M_s_ or P301L mutant in contrast to normal tau where it is relatively shielded. Thus, the dramatic sensitivity of P301L to seeds can be explained by an increased exposure of the aggregation-prone ^306^VQIVYK^311^ sequence. These data suggest that M_s_ functions catalytically to convert normal tau into aggregates. Thus, the proposed seeding mechanism of M_s_ may be generalized to tauopathies that are not caused by mutations.

### Perturbing metastable tau structures

Ensemble averaging methods, such as NMR, have had limited success in understanding the solution conformations of tau under physiological conditions. They have revealed secondary structure propensities of key regions and proposed the existence of local contacts^2,7,22,23,51^. However, capturing more transient or low population local conformations has been difficult. This is confounded by poor signal to noise, requiring long acquisition times at high concentrations, and non-physiological temperatures to suppress protein aggregation. As such, capturing transient but important local structural signatures have been challenging with classical structural biology methods. Both experiment and simulation have shown that weak local structure may play key roles in limiting aggregation of globular proteins during translation and that these structural elements may play even bigger roles in intrinsically disordered proteins^52,53^. Thus, local structures that bury proximal amyloid sequences may be a general evolutionary design principle that controls aggregation.

Our study has suggested that local structure encompassing the amyloid motif ^306^VQIVYK^311^ regulates aggregation of tau and that the P301L mutation increases susceptibility to conformational changes that expose the ^306^VQIVYK^311^ amyloid motif. While these differences are subtle, we observe that P301L-mediated structural rearrangements only manifest under moderate stress conditions (i.e. heat, seed). Hence, as compared to NMR, real-time assays, such as FRET, and XL-MS that kinetically trap conformations are more appropriate to detect metastable sub-populations. These data may explain the elusiveness of a biophysical basis of the cluster of pathogenic mutations near ^306^VQIVYK^311^.

Simulations predict that repeat interfaces could encode local structures that are compatible with a β-hairpin and that the P301L mutation, dramatically shifted the equilibrium away from collapsed hairpins to extended fibril-like conformations. Our findings are consistent with published NMR data -- PGGG sequences in tau can adopt type II β-turns^7^ and that the P301L mutation increases local β-strand propensity^27^. Thus, our work supports the structural and functional findings that metastable local structures in tau are destabilized by disease-associated mutations.

### Peptides capture local structure observed in tau

Guided by our simulations, we predicted that a local fragment spanning the interface between repeat 2 and 3 should encode a minimal structure necessary to replicate this aggregation phenomenon. We examined whether structural perturbations influenced aggregation propensity in a peptide model system that captures this local structural element. The WT tau interface peptide model containing ^306^VQIVYK^311^ did not aggregate spontaneously; however, single point substitutions of 6 disease-associated mutations immediately N-terminal to ^306^VQIVYK^311^ consistently induced spontaneous aggregation. Given that destabilization of local structure around ^306^VQIVYK^311^ promotes aggregation, stabilizing local structure should rationally mitigate aggregation. By promoting a β-hairpin structure *via* tryptophan zipper motifs or by using isoelectric forces, a P301L-containing tau peptide had an inhibited propensity to aggregate.

Our data support the hypothesis that local forces are key to preventing aggregation of tau by maintaining specific local structures. Tau is generally considered to be an intrinsically disordered protein, and therefore long-range contacts are unlikely to play a significant role in stability. Published NMR experiments support local structure formation of these regions in tau. Spectra of tau RD (K18; amino acids 244 – 372) overlaps with a N- and C-terminally expanded tau RD (K32; amino acids 198 – 394) and even with the splice isoform of tau RD missing repeat 2 (K19; amino acids 244-372 with 275-306 deleted)^7,51^, suggesting that adding residues and even deleting an entire repeat have minimal effects on the local structure. Thus, the conformations of local structures in tau are disproportionally more important to its properties compared with structured proteins. This suggests that peptide fragment models are a valid surrogate and can encapsulate the most relevant endogenous structural elements for investigating aggregation of tau.

### Tau mutations drive disease by promoting rapid aggregation

Disease-associated mutations found near tau’s amyloid motif, such as P301L or P301S, have no definitive biophysical mechanism but are nevertheless widely used in cell and animal models of disease^25,26^. Using our peptide, tau RD and FL-tau model systems, we observe that key mutations dramatically alter aggregation rates on similar time scales *in vitro* and seed in cell models. We therefore provide an explanation for the toxic-gain-of-function for several mutations.

Previous reports have studied intra-repeat interactions, with the assumption that each repeat functions independently within tau RD^33^. Their peptide models have shown a relationship between the length of a peptide fragment and the seeding capacity of tau to define repeat 3 as the minimal sequence necessary to act as a fully functional seed^33^. Our model defines a minimal regulatory sequence that limits spontaneous aggregation and suggests that inter-repeat structural elements modulate aggregation propensity. The composition of these inter-repeat sequences, governed by missense mutations, directly impacts the stability of local structures and aggregation propensity. It is tempting to speculate that local structure surrounding each of the four inter-repeat regions plays independent roles in the exposure of amyloid sequences. This modular nature of the tau RD region may explain how these independent regions can lead to distinct structures that define tau prion strains^4^. A more comprehensive structure-function analysis of other repeat interfaces may help explain how each inter-repeat element contributes to the formation of different tau structures.

### Splice variants modulate VQIVYK aggregation propensity

The expression levels of the two major isoform types of tau in the central nervous system – 3R and 4R – are similar in the adult brain^29^. However, the 3R:4R ratio of aggregate deposits is disproportionally shifted towards 4R in most tauopathies^30^. Mutations in tau that affect alternative splicing and generate excess 4R isoforms correlate with some genetic tauopathies^31,32^. The N-terminal flanking sequences leading into ^306^VQIVYK^311^ differ by four amino acids between the two isoforms. We find that these two isoforms have drastically different aggregation propensities in the presence of disease-associated mutations (t_1/2_ = 7 hours vs t_1/2_ > 100 hours, respectively). Chimeras of R1R3 / R2R3 transition from aggregation-resistant to aggregation-prone as they lose R1 N-terminal flanking character. The ability of an R1 leading strand to mitigate ^306^VQIVYK^311^ aggregation may explain why 4R tau correlates more closely with pathology. Thus, inter-repeat contacts may explain aggregation propensities of tau isoforms in disease. Encouraging data for a tau vaccine targeting a ^300^HXPGGG^304^ sequence suggests it’s possible to utilize inter-repeat regions to select between pathogenic and non-pathogenic conformations of tau^54^.

### *Cis*-*trans* isomerization and prolyl isomerases

Studying the missense mutations in tau has generated valuable disease models ^26,35^; however, the majority of human tauopathies have no observed genetic mutation in tau^34^. Critical proline residues N-terminal to the amyloid motif can isomerize into *cis* or *trans* rotamers spontaneously or through unidentified cellular mechanisms. We observe that P301 *cis* and *trans* rotamers have distinct aggregation propensities *in vitro*. In fact, the aggregation kinetics for a *trans* rotamer of P301 are on par with some well-defined disease mutants (N296Δ, V300I). The concept of proline isomerization triggering aggregation into amyloid is not novel, as this is an accepted mechanism of β2-microglobulin aggregation in kidney dialysis amyloidosis^55^. Other proline residues outside of the tau repeat domain have also been proposed to undergo proline isomerization^48^. Our proposed model suggests a possible mechanism whereby WT tau aggregation could be controlled *in vivo*: specific prolyl isomerization events – possibly triggered by cellular proline isomerases – could trigger spontaneous aggregation by modulating inter-repeat structural elements.

## Conclusions

We propose that sequences N-terminal to tau’s amyloid motif forms local contacts consistent with a β-hairpin-like compact structure. This shields the amyloid motif and mitigates aggregation (**Figure 8**). This represents a simple yet comprehensive model of tau aggregation that unifies key observations throughout tau literature.

**Figure 8.**
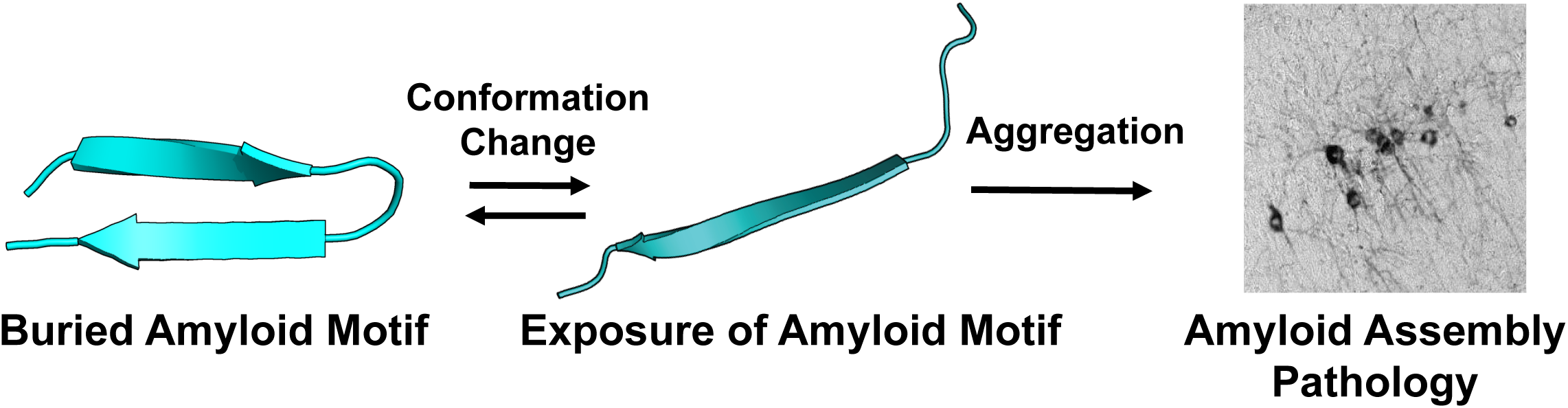
Molecular model of tau amyloid domain structural rearrangement and subsequent aggregation. Naïve tau monomer (left) exists with a propensity to form a relatively collapsed structure, which buries the amyloid domain ^306^VQIVYK^311^. In the presence of disease-associated mutations, proline isomerization events, or certain splice isoforms, the equilibrium is shifted to disfavor local compact structure. This exposes the aggregation prone ^306^VQIVYK^311^amyloid motif and enhances aggregation propensity, leading to subsequent tau pathology.

Algorithms that identify potential amyloid nucleating regions, such as TANGO, have indicated that nearly 75% of aggregation nucleating regions in the human proteome use two or more “gatekeeper” residues, with proline being the most common single gatekeeping residue^56^. These gatekeeping residues are more likely than average to be the site of disease-associated missense mutations and are consistent with our identification of gatekeeping residues near tau’s amyloid motif. Thus, local flanking sequences and their structural contacts may play an important role in mitigating aggregation propensity in tau and likely other intrinsically disordered proteins.

Finally, the identification and characterization of metastable compact structures encompassing ^306^VQIVYK^311^ may itself prove to be a valuable therapeutic target. One might be able to shift the structural rearrangement of tau amyloid motif from exposed (aggregation prone) to buried (inert) using small molecules, antibodies, or cellular co-factors. Our results indicate that subtle changes in local structure have immense functional ramifications; therefore, small molecules that shift this structural equilibrium modestly may have significant benefits.

## METHODS

### Recombinant Full-Length tau and tau RD Production

We utilized several forms of recombinant tau. Full-length (FL) wildtype and P301L tau were cloned into pet28b vector (a kind gift from Dr. David Eisenberg) using Ncol and Xhol sites. The plasmid was transformed into BL21-Gold (DE3) cells. Cells were grown in 1x Terrific Broth media to OD600 1.4 and induced with 1 mM IPTG for 3 hours at 37°C. The cells were harvested and lysed in 50 mM Tris, 500 mM NaCl, 1 mM β-Mercaptoethanol, 20 mM imidazole, 1 mM PMSF, pH 7.5, using an Omni Sonic Ruptor 400. The lysates were centrifuged, the supernatant was applied to a Ni-NTA column and eluted with 50 mM Tris, 250 mM NaCl, 1 mM β-Mercaptoethanol, 300 mM imidazole. Eluting fractions containing tau were desalted into 50 mM MES, 50 mM NaCl, 1mM β-Mercaptoethanol (pH 6.0) by PD-10 column GE. Exchanged fractions were applied to a HiTrap SP HP (GE) and eluted with a 50mM-1M NaCl gradient. Tau containing fractions were concentrated on an Amicon-15 concentrator and applied to a Superdex 200 Increase 10/300 GL (GE) and eluted into 1x PBS (136.5 mM NaCl, 2.7 mM KCl, 10 mM Na_2_HPO_4_, 1.8 mM KH_2_PO_4_, pH 7.4). Tau RD wildtype and mutants were expressed and purified as previously described ^57^. Aliquots were all stored at −80°C in 1x PBS. Tau seeding monomer (M_s_) was produced as previously described ^16^.

### ThT Fluorescence Aggregation Assays

Wildtype or mutant FL tau and tau RD protein was diluted in 1x PBS with 5µM β-Mercaptoethanol and boiled at 95°C for 5 min. A final concentration of 4.4µM heparin or 33nM M_s_ seed was added to 4.4µM tau or tau RD protein in a 60µL volume mixed with 25 µM ThT and aliquoted into a 96-well clear bottom plate. Peptides were disaggregated as previously described^58^, lyophilized, and resuspended in 2x PBS. 25 µM ThT was added to 200uL of 200 µM peptide in a 96-well clear bottom plate. All conditions were done in triplicates (except for the R2R3-IEZip experiment) at room temperature. ThT kinetic scans were run every 5 minutes on a Tecan M1000 plate reader at 446 nm Ex (5 nm bandwidth), 482 nm Em (5 nm bandwidth). Blank wells containing buffer and ThT were subtracted from experimental values. Samples producing signal to background (ThT only) with ratios only greater than 2:1 were considered and these values were normalized to the maximum amplitude in each condition. The data were plotted, and the t_1/2_ values were derived from a non-linear regression model fitting in GraphPad Prism.

### Transmission electron microscopy (TEM)

An aliquot of 5 μl of sample was placed onto a glow-discharged Formvar-coated 400-mesh copper grids for 30 seconds, washed with distilled water, and then negatively stained with 2% uranyl acetate for 1 min. Images were acquired on a Tecnai G^2^ spirit transmission electron microscope (FEI, Hillsboro, OR), serial number: D1067, equipped with a LaB_6_ source at 120kV using a Gatan ultrascan CCD camera.

### Tau Biosensor Cells

Biosensor cells were plated into 96-well plates at 20,000 cells per well. For tau and tau RD experiments, after 5 days of incubation with heparin or M_s_, 10µL of 4.4µM aggregated protein material was mixed with 1.25µL lipofectamine and 8.75µL Opti-MEM, incubated at room temperature for 30 min, and added to cell media. The “t=0” samples were prepared in the same way but straight from the freezer aliquots. After 2 days, cells were harvested with 0.05% trypsin, then resuspended in Flow buffer (1x HBSS, 1% FBS, 1 mM EDTA, 1x DPBS) and analyzed by flow cytometry. For peptide experiments, 10 µg of aggregated peptide material was added to 0.5 uL lipofectamine and Opti-MEM to a total volume of 10 µL, incubated at room temperature for 30 min, and added directly to cell media. After 3 days, cells were harvested with 0.05% trypsin, then resuspended in Flow buffer and analyzed by flow cytometry. All conditions were done in triplicates. The Trp Zip biosensor cells expressing the tryptophan zipper motifs flanking the R2R3 element in tau RD were generated as previously described^25^.

### Tau intramolecular FRET

10-fold excess TCEP was added to 100 µM tau RD, the atmosphere was deoxygenated with N_2_ gas, and incubated at RT for 30 minutes. To label, C_5_ maleimide Alexa-488 and C_2_ maleimide Alexa-647 were dissolved in DMSO and added in 5-fold excess to tau RD, and incubated at 4°C overnight. Alexa-labelled tau RD was purified using an 8 kDa cut-off mini dialysis exchanged into 1x PBS overnight. FRET measurements were carried out at 100nM tau RD labeled protein samples. Mi and Ms were purified as described above. For M_i_ and M_s_ intramolecular FRET experiments, 4uM protein samples were reduced with 10-fold excess TCEP, the atmosphere was deoxygenated with N_2_ gas, and incubated at RT for 30 minutes. To label, C_5_ maleimide Alexa-488 and C_2_ maleimide Alexa-647 were dissolved in DMSO and added in 5-fold excess to M_i_ and M_s_, and incubated at 4°C overnight. Alexa-labelled M_i_ and M_s_ was purified using an 8 kDa cut-off mini dialysis exchanged into 1x PBS overnight. FRET measurements were carried out at 100nM M_i_ and M_s_ labeled protein samples. FRET efficiencies were calculated as a function of acceptor emission, given E = (I_AD_ε_AA_-I_AD_ε_AD_)/I_AA_ε_DD_ where I_AA_ is Acceptor intensity following Acceptor excitation, I_AD_ is Acceptor intensity following Donor excitation, and ε is the extinction coefficient of the fluorophores at given excitations and emissions. Alexa 488 Ex/Em was measured at 490 nm/520 nm, Alexa 647 Ex/Em 647 nm/670 nm with a 5 nm bandwidth for both. As a control, spectra for free 100nM Alexa 647 and Alexa 488 were acquired at 37°C, 50°C and 75°C at two time points (t=0 and t=30 minutes). FRET efficiencies remained flat and were not perturbed by heating (**Supplemental Table 2**).

### Crosslinking, sample processing and LC-MS/MS analysis

Preparation of tau RD was crosslinked at a total protein concentration of 1.0 mg/mL using 100 µg of starting material. The crosslinking buffer was 1X PBS with 3mM DTT. Five replicates for each condition (37°C, 50°C, and 75°C) were prepared. Samples for 50°C and 75°C conditions were equilibrated at the appropriate temperature for 1 hour before crosslinking. The crosslinking reaction was initiated by adding disuccinimidyl suberate (DSS) stock solution (25 mM DSS-d_0_ and –d_12_, Creative Molecules) in DMF to a final concentration of 1 mM. Samples were further incubated at 37°C, 50°C, or 75°C for 1 min with 350 RPM shaking. Excess reagent was quenched by addition of Tris (pH 7.5) to 100 mM and incubation at 37°C for 30 min, and subsequently flash frozen by liquid nitrogen and evaporated to dryness by lyophilization. Proteins were resuspended in 8 M urea, reduced with 2.5 mM TCEP (37°C, 30 min) and alkylated with 5mM iodoacetamide (30min, room temperature, protected from light). The sample solutions were diluted to 1 M urea with 50 mM ammonium hydrogen carbonate and trypsin (Promega) was added at an enzyme-to-substrate ratio of 1:50. Proteolysis was carried out at 37°C overnight followed by acidification with formic acid to 2% (v/v). Samples were then purified by solid-phase extraction using Sep-Pak tC18 cartridges (Waters) according to standard protocols. Samples were evaporated to dryness and reconstituted in water/acetonitrile/formic acid (95:5:0.1, v/v/v) to a final concentration of approximately 0.5 µg/µl. 2µL each were injected for duplicate LC-MS/MS analyses on an Eksigent 1D-NanoLC-Ultra HPLC system coupled to a Thermo Orbitrap Fusion Tribrid system. Peptides were separated on self-packed New Objective PicoFrit columns (11cm x 0.075mm I.D.) containing Magic C_18_ material (Michrom, 3µm particle size, 200Å pore size) at a flow rate of 300nL/min using the following gradient. 0-5min = 5 %B, 5-95min = 5-35 %B, 95-97min = 35-95 %B and 97-107min = 95 %B, where A = (water/acetonitrile/formic acid, 97:3:0.1) and B = (acetonitrile/water/formic acid, 97:3:0.1). The mass spectrometer was operated in data-dependent mode by selecting the five most abundant precursor ions (m/z 350-1600, charge state 3+ and above) from a preview scan and subjecting them to collision-induced dissociation (normalized collision energy = 35%, 30ms activation). Fragment ions were detected at low resolution in the linear ion trap. Dynamic exclusion was enabled (repeat count 1, exclusion duration 30sec).

### Analysis of mass spectrometry data

Thermo.raw files were converted to the open.mzXML format using msconvert (proteowizard.sourceforge.net) and analyzed using an in-house version of xQuest^50^. Spectral pairs with a precursor mass difference of 12.075321 Da were extracted and searched against the respective FASTA databases containing tau (TAU_HUMAN P10636-8) or with a P301L/S substitution. xQuest settings were as follows: Maximum number of missed cleavages (excluding the crosslinking site) = 2, peptide length = 5-50 aa, fixed modifications = carbamidomethyl-Cys (mass shift = 57.021460 Da), mass shift of the light crosslinker = 138.068080 Da, mass shift of mono-links = 156.078644 and 155.096428 Da, MS^1^ tolerance = 10 ppm, MS^2^ tolerance = 0.2 Da for common ions and 0.3 Da for crosslink ions, search in ion-tag mode. Post-search manual validation and filtering was performed using the following criteria: xQuest score > 25, mass error between −2.2 and +3.8ppm, %TIC > 10, and a minimum peptide length of six aa. In addition, at least four assigned fragment ions (or at least three contiguous fragments) were required on each of the two peptides in a crosslink. False discovery rates (FDR’s) for the identified crosslinks were estimated using xprophet^59^ and estimated to be 1.3 - 10% (**Supplemental Figure 4, Supplemental Table 3**). At each temperature, the 5 replicate datasets were compared and only crosslinks present in 5 of the 5 datasets were used to generate a consensus dataset (**Supplemental Table 4**). Crosslink data with information of crosslinked residue positions and nseen (frequency) was visualized using customized gnuplot scripts.

### Model generation of tau RD using ROSETTA

The backbone NH, N, CA, CB and C=O chemical shift assignments for the tau fragment from 243-368 (bmrbId=19253) were used in CS-Rosetta to generate fragment libraries for subsequent model refinement ^60^. First, chemical shift parameters were used to predict backbone torsional angles using TALOS to generate a CS-guided fragment library representing the conformations of the protein^60^. For the *ab initio* ROSETTA calculations, the tau RD sequence was used to generate 3-mer and 9-mer fragments derived from the protein data bank using the fragment picker tool^40^. The Rosetta energy function was used to assemble and iteratively refine 5000 structural models using each set of fragments^40,41,61^. Cα–based root mean square deviations were computed in Rosetta for tau RD and hairpin fragments. Clustering analysis of the tau RD ensemble showed similar results yielding median root mean square deviations of 19.5 Ang^2^ and 19.75 Ang^2^ for *ab initio* and CS-ROSETTA simulations, respectively (data not shown). Ensemble wide calculation of cα-cα end to end distances between residues 264-280, 295-311, 327-343 and 359-375 were carried out using a python script. All simulations were performed on UTSW’s biohpc computing cluster. All plots were generated with gnuplot. Images were created using Pymol.

### Peptide Synthesis

All peptides were synthesized as ordered by Genscript with N-terminal acetylation and C-terminal amidation modifications. Peptides were purified to >95% purity by FPLC *via* an Agilent ZORBAX StableBond 250 mm C8 column.

### Flow Cytometry

A BD LSRFortessa was used to perform FRET flow cytometry. To measure CFP and FRET, cells were excited with the 405 nm laser, and fluorescence was captured with a 405/50 nm and 525/50 nm filter, respectively. To measure YFP, cells were excited with a 488 laser and fluorescence was captured with a 525/50 nm filter. To quantify FRET, we used a gating strategy where CFP bleed-through into the YFP and FRET channels was compensated using FlowJo analysis software. We then created a plot of FRET vs. CFP and introduced a triangular gate to assess the number of FRET-positive cells, as previously described^16^. For each experiment, at least 10,000 single cells per replicate were analyzed. Data analysis was performed using FlowJo v10 software (Treestar).

### MD Simulations

Well-Tempered Metadynamics^62^ was employed to enable accelerated conformational sampling and to construct the associated free energy surface. Metadynamics was performed on a two-dimensional space of parallel-β sheet content and anti-parallel sheet content. To increase search efficiency in oligomeric space, we have incorporated conformational symmetry constraints, which have been shown to enable sampling of multi-polymer landscapes^43^. The initial dodecahedron simulation box was constructed from a trimer of a randomly unfolded structure of 295-311 by adding 7587 SPCE explicit waters and 3 neutralizing Cl ions (one for each monomer). The AMBER99sb-ildn force-field^63^ was used for all simulations. After an initial 1009 steepest descent steps of converged energy minimization, 10 ns of NVT and 20 ns of NPT (first 10 with Berendsen^64^ and the last 10 with Parrinello-Rahman^65^ barostats) equilibrations were performed. The subsequent production level trajectories are based on 5 fs time steps using hydrogen-only virtual sites^66^. Production level trajectories were obtained for an NPT ensemble with Parrinello-Rahman barostat, and periodic boundary conditions with Particle Mesh Ewald (PME) ^67^ summation for long-range electrostatics. The tuned well-tempered metadynamics parameters are 10, 1.4 kJ/mole, and 0.3 for bias factor, Gaussian height, collective variable space Gaussian widths, respectively. The Gaussian perturbations were included into MD every 2.5 ps using the PLUMED package^68^ as an external patch to Gromacs-5.0.4^69^. A total of 18 μs trajectories were generated, 9 μs for wild type and 9 μs for the P301L mutant, over a total of 6 independent runs. All simulations were done on UTSW’s bioHPC computing cluster.

### Statistics

All statistics were calculated using GraphPad Prism 8.0. Three independent ThT experiments were run for each condition. The data were normalized to the highest amplitude and averages and standard deviations were plotted. Plots were fitted to a non-linear regression model, from which t_1/2_ values were derived. t_1/2_ error represents a 95% CI. Flow cytometry cell aggregation was conducted in 3 independent experiments, whose values are plotted. Error bars represent a 95% CI.

### Data Availability

ThT data for tau, tau RD and peptide experiments is available in Supplemental Table 1. Raw crosslinking mass spectrometry data is available in Supplemental Table 3 and 4. Other supporting data available upon request from the authors.

## Acknowledgements

This work was supported by grants from the Tau Consortium and NIH grants awarded to 1R01NS071835 (M.I.D.), the Effie Marie Cain Endowed Scholarship (L.A.J.) and the Heising-Simons Foundation (M.M.L.). We appreciate the help of the Structural Biology Laboratory and Proteomics Core Facility at the University of Texas Southwestern Medical Center.

## Author Contributions

K.D. M.I.D., and L.A.J. conceived and designed the overall study. K.D. performed *in vitro* peptide assays, intramolecular FRET, flow cytometry, and cell models. D.C. performed *in vitro* protein assays, cell models, crosslink mass spectrometry, and ROSETTA simulations. Z.H. developed the tau *in vitro* seeded aggregation assays. O.M.K. assisted with tau and tau RD purifications. V.A.P. helped with cell seeding assays. B.D.R. and D.R.W. performed electron microscopy. L.S. and M.M.L. performed molecular dynamics simulations. K.D., D.C. and L.A.J. wrote the manuscript, and all authors contributed to its improvement.

## References

1. Cleveland, D. W., Hwo, S. Y. & Kirschner, M. W. Physical and chemical properties of purified tau factor and the role of tau in microtubule assembly. Journal of Molecular Biology 116, 227–247 (1977).

2. Eliezer, D. et al. Residual structure in the repeat domain of tau: Echoes of microtubule binding and paired helical filament formation. Biochemistry 44, 1026–1036 (2005).

3. Fitzpatrick, A. W. P. et al. Cryo-EM structures of tau filaments from Alzheimer’s disease. Nature 56, 343 (2017).

4. Sanders, D. W. et al. Distinct Tau Prion Strains Propagate in Cells and Mice and Define Different Tauopathies. Neuron 82, 1271–1288 (2014).

5. Sawaya, M. R. et al. Atomic structures of amyloid cross-beta spines reveal varied steric zippers. Nature 447, 453–457 (2007).

6. Bergen, von, M. et al. Mutations of tau protein in frontotemporal dementia promote aggregation of paired helical filaments by enhancing local beta-structure. J. Biol. Chem. 276, 48165–48174 (2001).

7. Mukrasch, M. D. et al. Sites of tau important for aggregation populate {beta}-structure and bind to microtubules and polyanions. Journal of Biological Chemistry 280, 24978–24986 (2005).

8. Falcon, B., Zhang, W., Murzin, A. G., Murshudov, G. & Holly, J. Structures of filaments from Pick’s disease reveal a novel tau protein fold. (2018).

9. Zhang, X. et al. RNA Stores Tau Reversibly in Complex Coacervates. bioRxiv 111245 (2017). doi:10.1101/111245

10. Ismail, T. & Kanapathipillai, M. Effect of cellular polyanion mimetics on tau peptide aggregation. J. Pept. Sci. 24, e3125 (2018).

11. Kuret, J. et al. Evaluating triggers and enhancers of tau fibrillization. Microsc. Res. Tech. 67, 141–155 (2005).

12. Zhao, J. et al. Glycan Determinants of Heparin-Tau Interaction. Biophys. J. 112, 921–932 (2017).

13. Kellogg, E. H. et al. Near-atomic model of microtubule-tau interactions. 1780, 1–9 (2018).

14. Mok, S.-A. et al. Mapping interactions with the chaperone network reveals factors that protect against tau aggregation. Nat Struct Mol Biol 25, 384–393 (2018).

15. Baughman, H. E. R., Clouser, A. F., Klevit, R. E. & Nath, A. HspB1 and Hsc70 chaperones engage distinct tau species and have different inhibitory effects on amyloid formation. J. Biol. Chem. 293, 2687–2700 (2018).

16. Mirbaha, H. et al. Inert and seed-competent tau monomers suggest structural origins of aggregation. eLife 7, (2018).

17. Sharma, A. M., Thomas, T. L., Woodard, D. R., Kashmer, O. M. & Diamond, M. I. Tau monomer encodes strains. eLife 7, (2018).

18. Elbaum-Garfinkle, S. & Rhoades, E. Identification of an aggregation-prone structure of tau. J. Am. Chem. Soc. 134, 16607–16613 (2012).

19. Chirita, C. N., Congdon, E. E., Yin, H. & Kuret, J. Triggers of full-length tau aggregation: a role for partially folded intermediates. Biochemistry 44, 5862–5872 (2005).

20. Walker, S., Ullman, O. & Stultz, C. M. Using intramolecular disulfide bonds in tau protein to deduce structural features of aggregation-resistant conformations. J. Biol. Chem. 287, 9591–9600 (2012).

21. Kadavath, H. et al. Folding of the Tau Protein on Microtubules. Angew. Chem. Int. Ed. Engl. 54, 10347–10351 (2015).

22. Mukrasch, M. D. et al. Highly populated turn conformations in natively unfolded tau protein identified from residual dipolar couplings and molecular simulation. J. Am. Chem. Soc. 129, 5235–5243 (2007).

23. Mukrasch, M. D. et al. Structural polymorphism of 441-residue tau at single residue resolution. PLoS Biol 7, e34 (2009).

24. Mutations. (2017).

25. Holmes, B. B. et al. Proteopathic tau seeding predicts tauopathy in vivo. Proc Natl Acad Sci USA 111, E4376–E4385 (2014).

26. Yoshiyama, Y. et al. Synapse loss and microglial activation precede tangles in a P301S tauopathy mouse model. Neuron 53, 337–351 (2007).

27. Fischer, D. et al. Structural and microtubule binding properties of tau mutants of frontotemporal dementias. Biochemistry 46, 2574–2582 (2007).

28. Karamanos, T. K., Kalverda, A. P., Thompson, G. S. & Radford, S. E. Mechanisms of amyloid formation revealed by solution NMR. Prog Nucl Magn Reson Spectrosc 88-89, 86–104 (2015).

29. Goedert, M., Wischik, C. M., Crowther, R. A., Walker, J. E. & Klug, A. Cloning and sequencing of the cDNA encoding a core protein of the paired helical filament of Alzheimer disease: identification as the microtubule-associated protein tau. Proc Natl Acad Sci USA 85, 4051–4055 (1988).

30. Williams, D. R. Tauopathies: Classification and clinical update on neurodegenerative diseases associated with microtubule-associated protein tau. Internal Medicine Journal 36, 652–660 (2006).

31. Schoch, K. M. M. et al. Increased 4R-Tau Induces Pathological Changes in a Human-Tau Mouse Model. Neuron 90, 941–947 (2016).

32. Hutton, M. et al. Association of missense and 5’-splice-site mutations in tau with the inherited dementia FTDP-17. Nature 393, 702–705 (1998).

33. Stöhr, J. et al. A 31-residue peptide induces aggregation of tau’s microtubule-binding region in cells. Nature Chemistry 9, 874–881 (2017).

34. Rizzu, P. et al. High Prevalence of Mutations in the Microtubule-Associated Protein Tau in a Population Study of Frontotemporal Dementia in the Netherlands. The American Journal of Human Genetics 64, 414–421 (1999).

35. Mocanu, M.-M. et al. The potential for beta-structure in the repeat domain of tau protein determines aggregation, synaptic decay, neuronal loss, and coassembly with endogenous Tau in inducible mouse models of tauopathy. J. Neurosci. 28, 737–748 (2008).

36. Furman, J. L., Holmes, B. B. & Diamond, M. I. Sensitive Detection of Proteopathic Seeding Activity with FRET Flow Cytometry. J Vis Exp e53205–e53205 (2015). doi:10.3791/53205

37. Leitner, A. et al. The Molecular Architecture of the Eukaryotic Chaperonin TRiC/CCT. Structure/Folding and Design 20, 814–825 (2012).

38. Joachimiak, L. A., Walzthoeni, T., Liu, C. W., Aebersold, R. & Frydman, J. The Structural Basis of Substrate Recognition by the Eukaryotic Chaperonin TRiC/CCT. Cell 159, 1042–1055 (2014).

39. Rinner, O. et al. Identification of cross-linked peptides from large sequence databases. Nat Meth 5, 315–318 (2008).

40. Ovchinnikov, S., Park, H., Kim, D. E., DiMaio, F. & Baker, D. Protein structure prediction using Rosetta in CASP12. Proteins 86 Suppl 1, 113–121 (2018).

41. Lange, O. F. et al. Determination of solution structures of proteins up to 40 kDa using CS-Rosetta with sparse NMR data from deuterated samples. Proc. Natl. Acad. Sci. U.S.A. 109, 10873–10878 (2012).

42. Mylonas, E. et al. Domain conformation of tau protein studied by solution small-angle X-ray scattering. Biochemistry 47, 10345–10353 (2008).

43. Lin, M. M. Leveraging symmetry to predict self-assembly of multiple polymers. Chemical Physics Letters 683, 347–351 (2017).

44. Kaufman, S. K. et al. Tau Prion Strains Dictate Patterns of Cell Pathology, Progression Rate, and Regional Vulnerability In Vivo. Neuron 92, 796–812 (2016).

45. Kier, B. L., Shu, I., Eidenschink, L. A. & Andersen, N. H. Stabilizing capping motif for beta-hairpins and sheets. Proc. Natl. Acad. Sci. U.S.A. 107, 10466–10471 (2010).

46. Vogel, M., Bukau, B. & Mayer, M. P. Allosteric regulation of Hsp70 chaperones by a proline switch. Molecular Cell 21, 359–367 (2006).

47. Gustafson, C. L. et al. A Slow Conformational Switch in the BMAL1 Transactivation Domain Modulates Circadian Rhythms. Molecular Cell 66, 447–457.e7 (2017).

48. Pastorino, L. et al. The prolyl isomerase Pin1 regulates amyloid precursor protein processing and amyloid-beta production. Nature 440, 528–534 (2006).

49. Torbeev, V. Y. & Hilvert, D. Both the cis-trans equilibrium and isomerization dynamics of a single proline amide modulate 2-microglobulin amyloid assembly. Proc. Natl. Acad. Sci. U.S.A. 110, 20051–20056 (2013).

50. Lim, J. et al. Pin1 has opposite effects on wild-type and P301L tau stability and tauopathy. Journal of Clinical Investigation 118, 1877–1889 (2008).

51. Mukrasch, M. D. et al. The ‘jaws’ of the tau-microtubule interaction. Journal of Biological Chemistry 282, 12230–12239 (2007).

52. Meng, W., Lyle, N., Luan, B., Raleigh, D. P. & Pappu, R. V. Experiments and simulations show how long-range contacts can form in expanded unfolded proteins with negligible secondary structure. Proc. Natl. Acad. Sci. U.S.A. 110, 2123–2128 (2013).

53. Ding, F., Jha, R. K. & Dokholyan, N. V. Scaling behavior and structure of denatured proteins. Structure/Folding and Design 13, 1047–1054 (2005).

54. Kontsekova, E., Zilka, N., Kovacech, B., Skrabana, R. & Novak, M. Identification of structural determinants on tau protein essential for its pathological function: novel therapeutic target for tau immunotherapy in Alzheimer’s disease. Alzheimers Res Ther 6, 45 (2014).

55. Eichner, T. & Radford, S. E. A Generic Mechanism of B2-Microglobulin Amyloid Assembly at Neutral pH Involving a Specific Proline Switch. Journal of Molecular Biology 386, 1312–1326 (2009).

56. Reumers, J., Maurer-Stroh, S., Schymkowitz, J. & Rousseau, F. D. Protein sequences encode safeguards against aggregation. Human Mutation 30, 431–437 (2009).

57. Frost, B., Ollesch, J., Wille, H. & Diamond, M. I. Conformational diversity of wild-type Tau fibrils specified by templated conformation change. Journal of Biological Chemistry 284, 3546–3551 (2009).

58. O’Nuallain, B. et al. Kinetics and Thermodynamics of Amyloid Assembly Using a High-Performance Liquid Chromatography-Based Sedimentation Assay. Meth. Enzymol. 413, 34–74 (2006).

59. Walzthoeni, T. et al. False discovery rate estimation for cross-linked peptides identified by mass spectrometry. Nat Meth 9, 901–903 (2012).

60. Shen, Y., Delaglio, F., Cornilescu, G. & Bax, A. TALOS+: A hybrid method for predicting protein backbone torsion angles from NMR chemical shifts. Journal of Biomolecular NMR 44, 213–223 (2009).

61. Raman, S. et al. NMR structure determination for larger proteins using backbone-only data. Science 327, 1014–1018 (2010).

62. Barducci, A., Bussi, G. & Parrinello, M. Well-tempered metadynamics: A smoothly converging and tunable free-energy method. Physical Review Letters 100, (2008).

63. Lindorff-Larsen, K. et al. Improved side-chain torsion potentials for the Amber ff99SB protein force field. Proteins 78, 1950–1958 (2010).

64. Berendsen, H. J. C., Postma, J. P. M., Van Gunsteren, W. F., Dinola, A. & Haak, J. R. Molecular dynamics with coupling to an external bath. J. Chem. Phys. 81, 3684–3690 (1984).

65. Parrinello, M. & Rahman, A. Polymorphic transitions in single crystals: A new molecular dynamics method. Journal of Applied Physics 52, 7182–7190 (1981).

66. Feenstra, K. A., Hess, B. & Berendsen, H. J. C. Improving efficiency of large time-scale molecular dynamics simulations of hydrogen-rich systems. Journal of Computational Chemistry 20, 786–798 (1999).

67. Darden, T., York, D. & Pedersen, L. Particle mesh Ewald: An N·log(N) method for Ewald sums in large systems. J. Chem. Phys. 98, 10089–10092 (1993).

68. Tribello, G. A., Bonomi, M., Branduardi, D., Camilloni, C. & Bussi, G. PLUMED 2: New feathers for an old bird. Computer Physics Communications 185, 604–613 (2014).

69. Abraham, M. J. et al. Gromacs: High performance molecular simulations through multi-level parallelism from laptops to supercomputers. SoftwareX 1-2, 19–25 (2015).

