## Supplemental Figures for "Tau local structure shields amyloid motif and controls aggregation propensity"

### Supplemental Figure 1

a

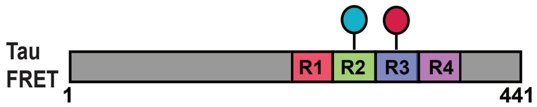


b

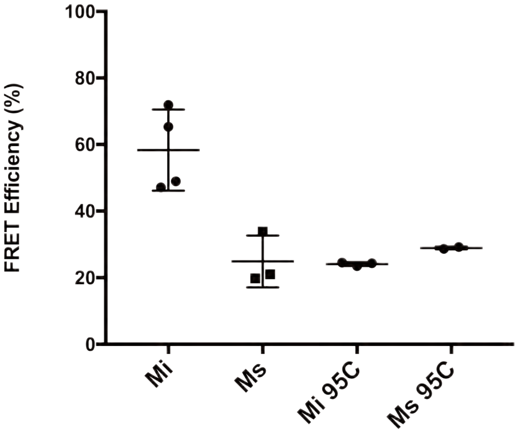

### Supplemental Figure 2

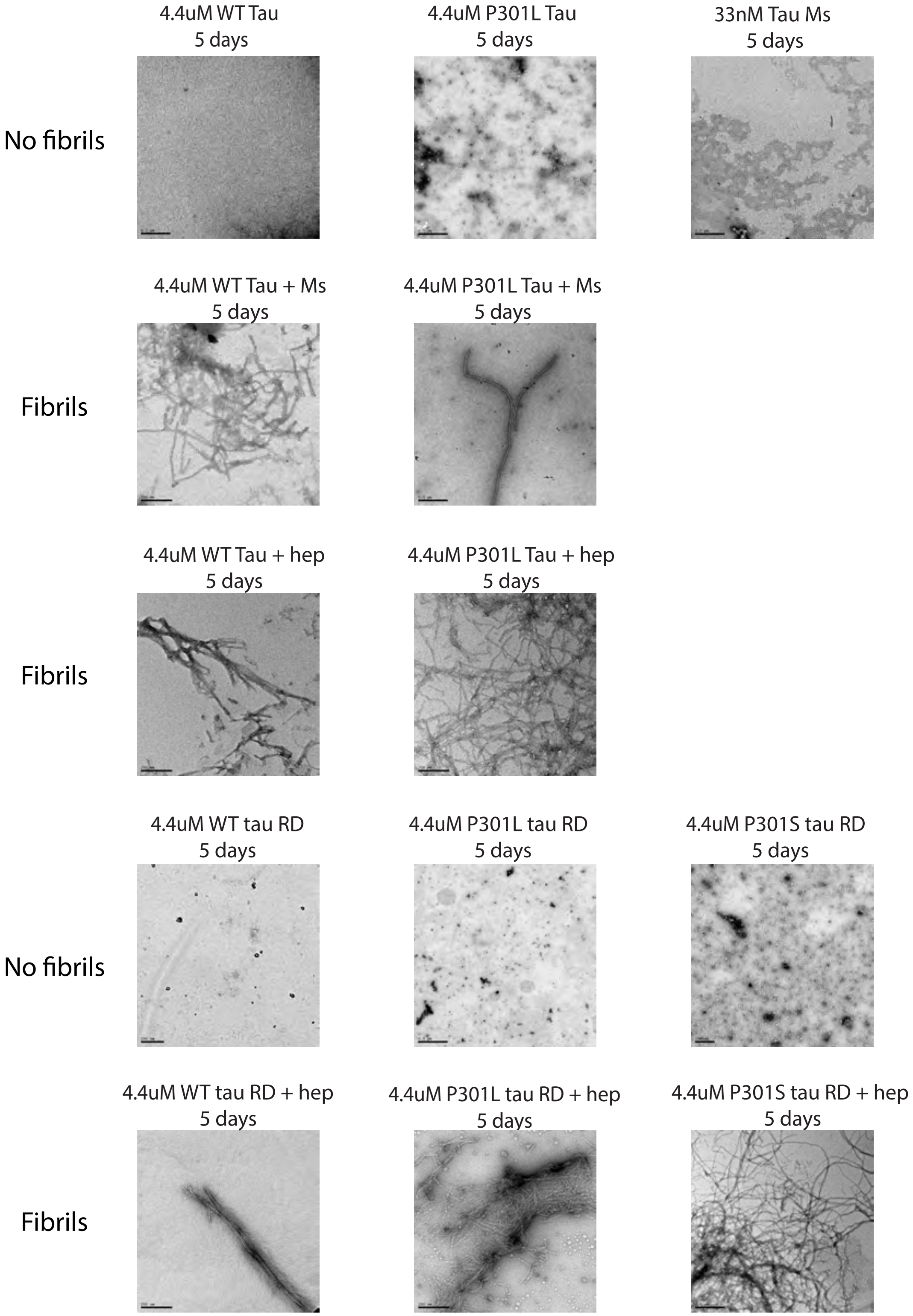

### Supplemental Figure 3

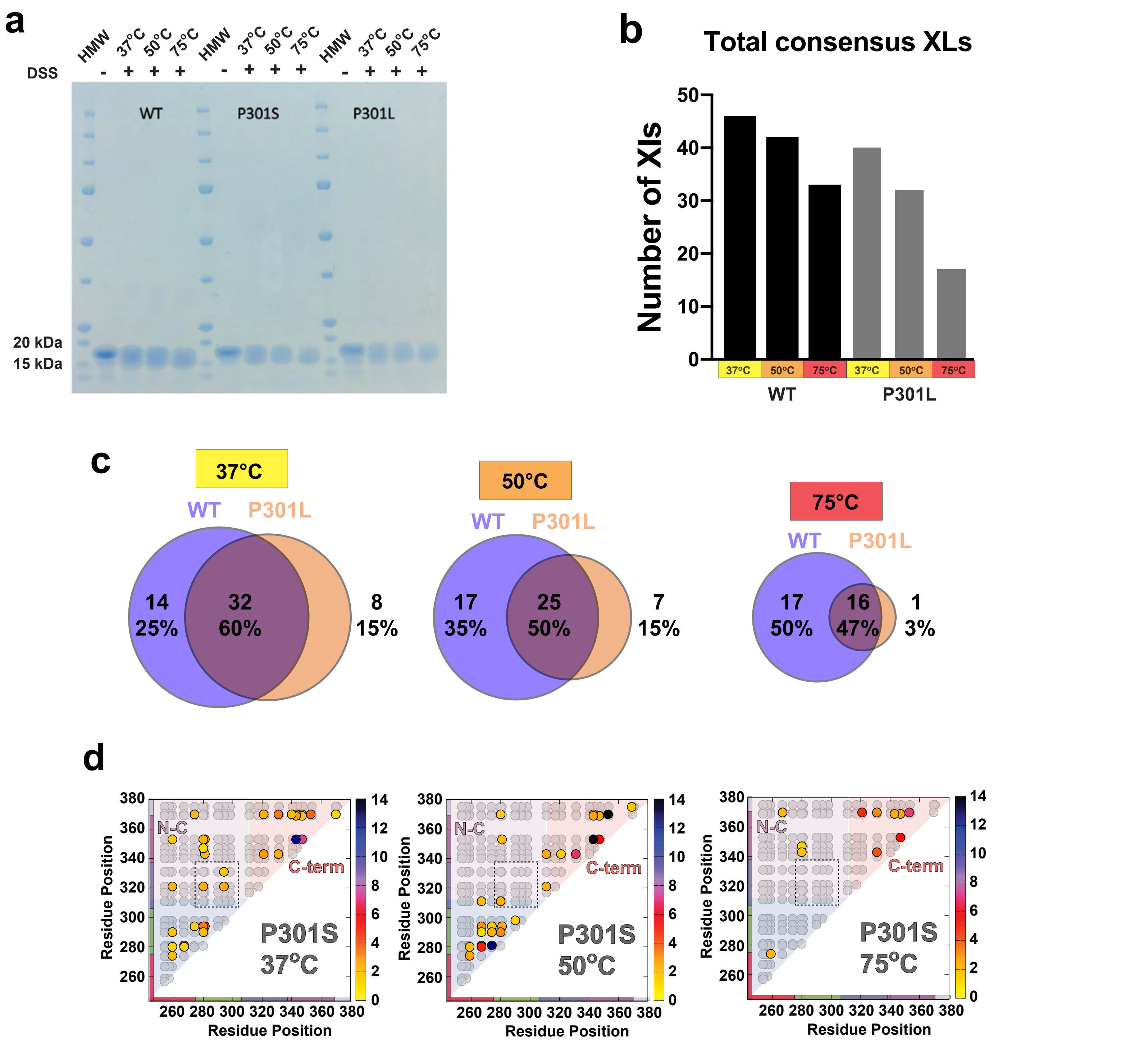

### Supplemental Figure 4

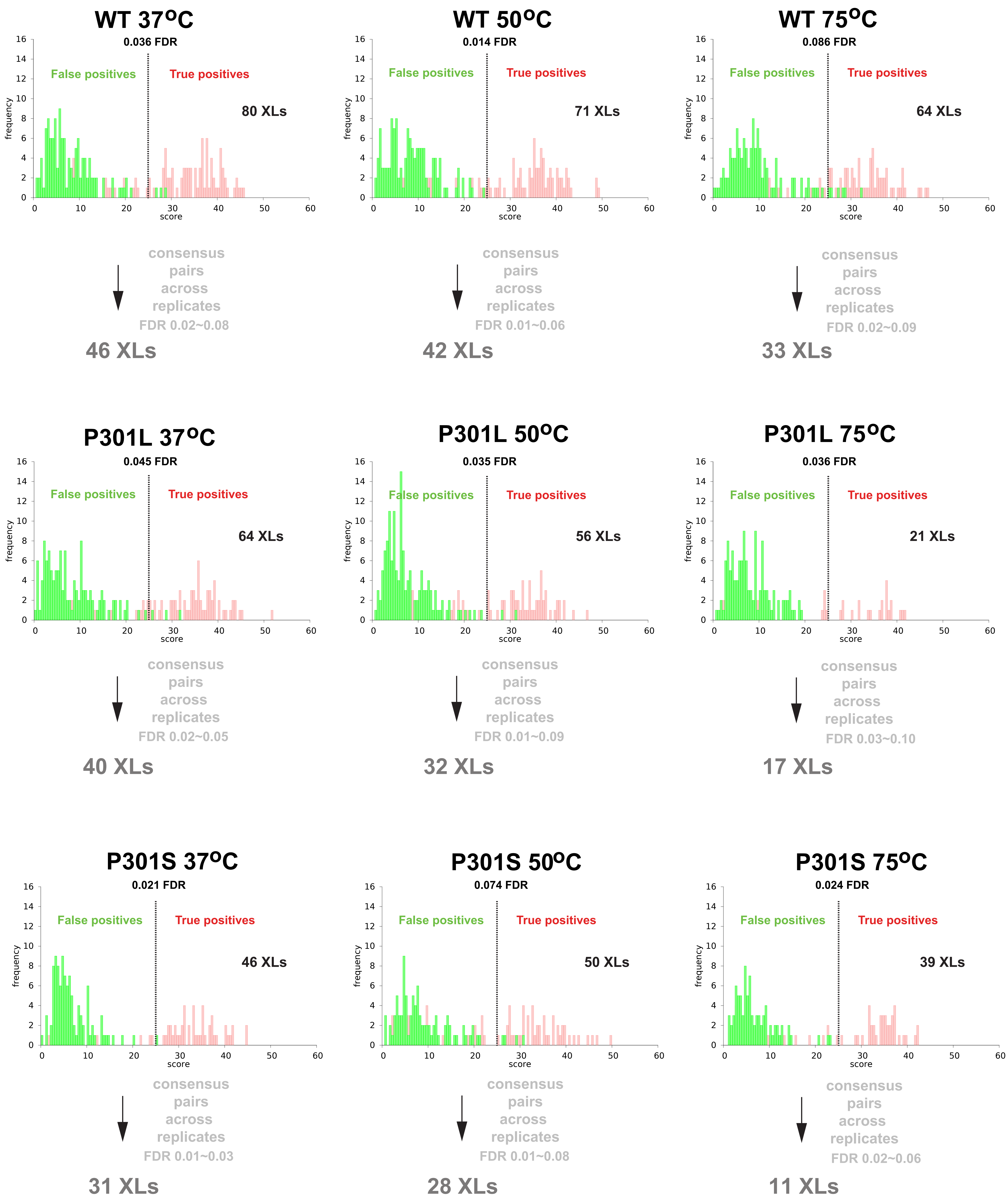

### Supplemental Figure 5

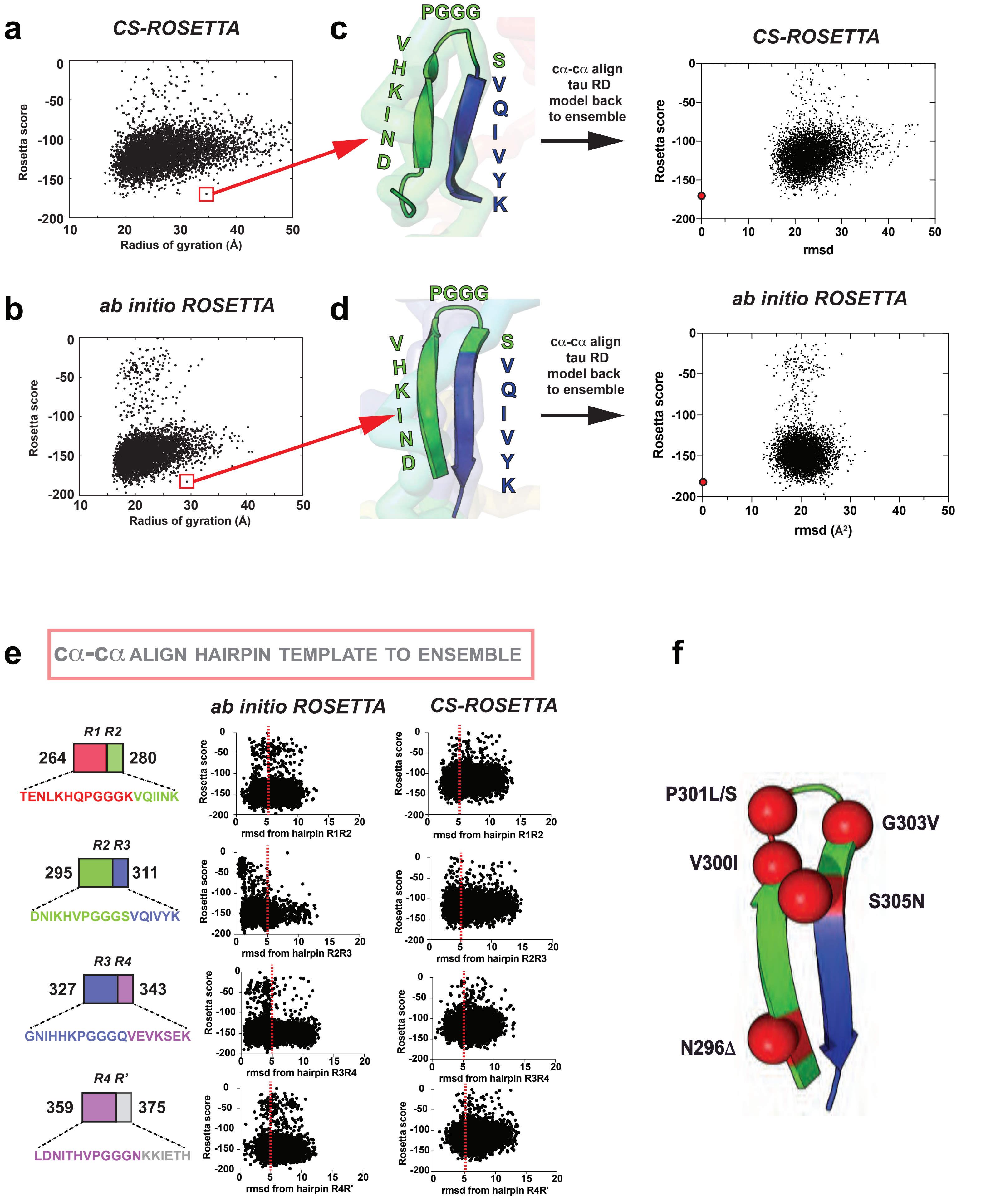

### Supplemental Figure 6

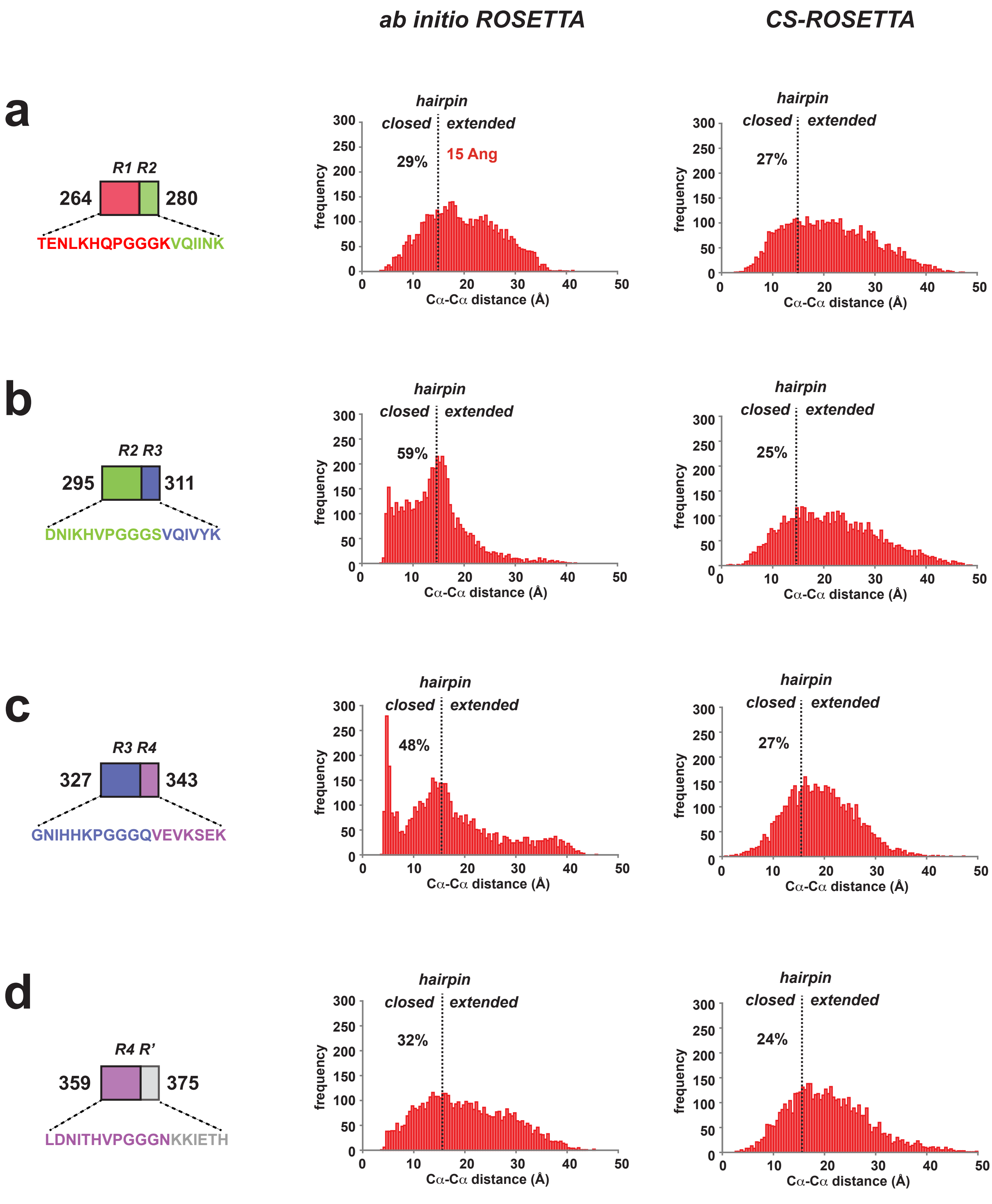

### Supplemental Figure 7

**a**

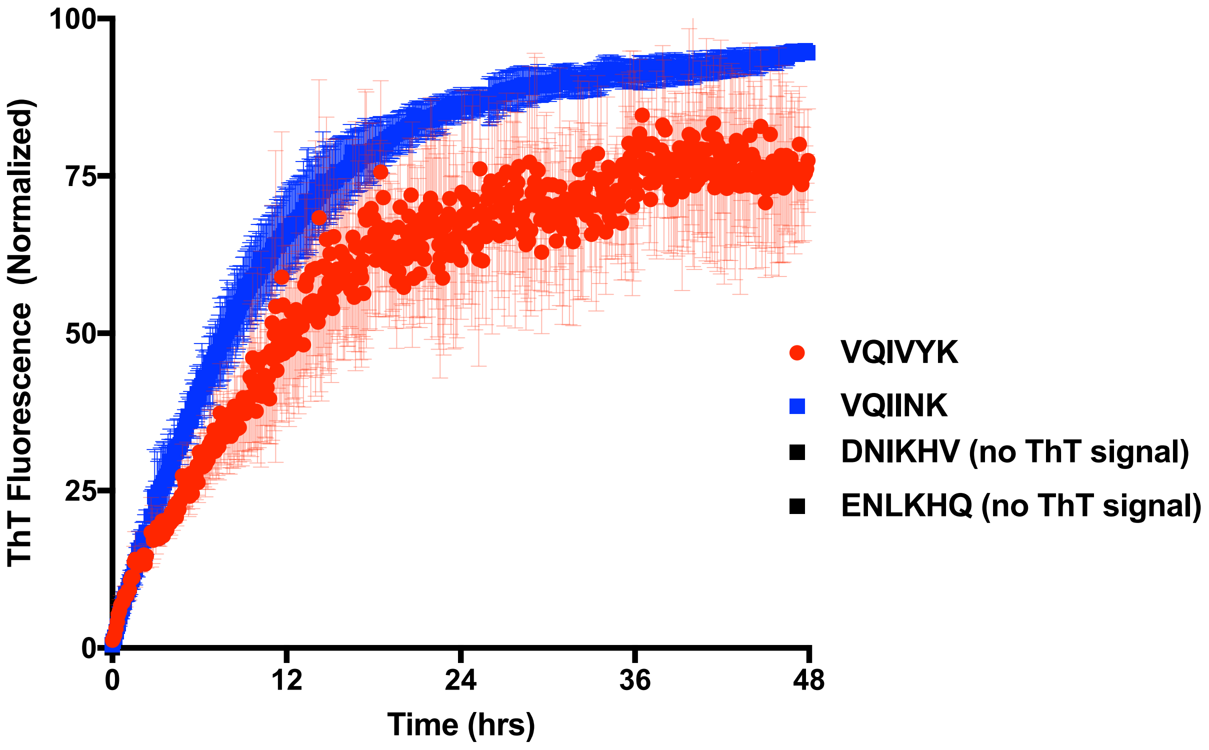

### Supplemental Figure 8

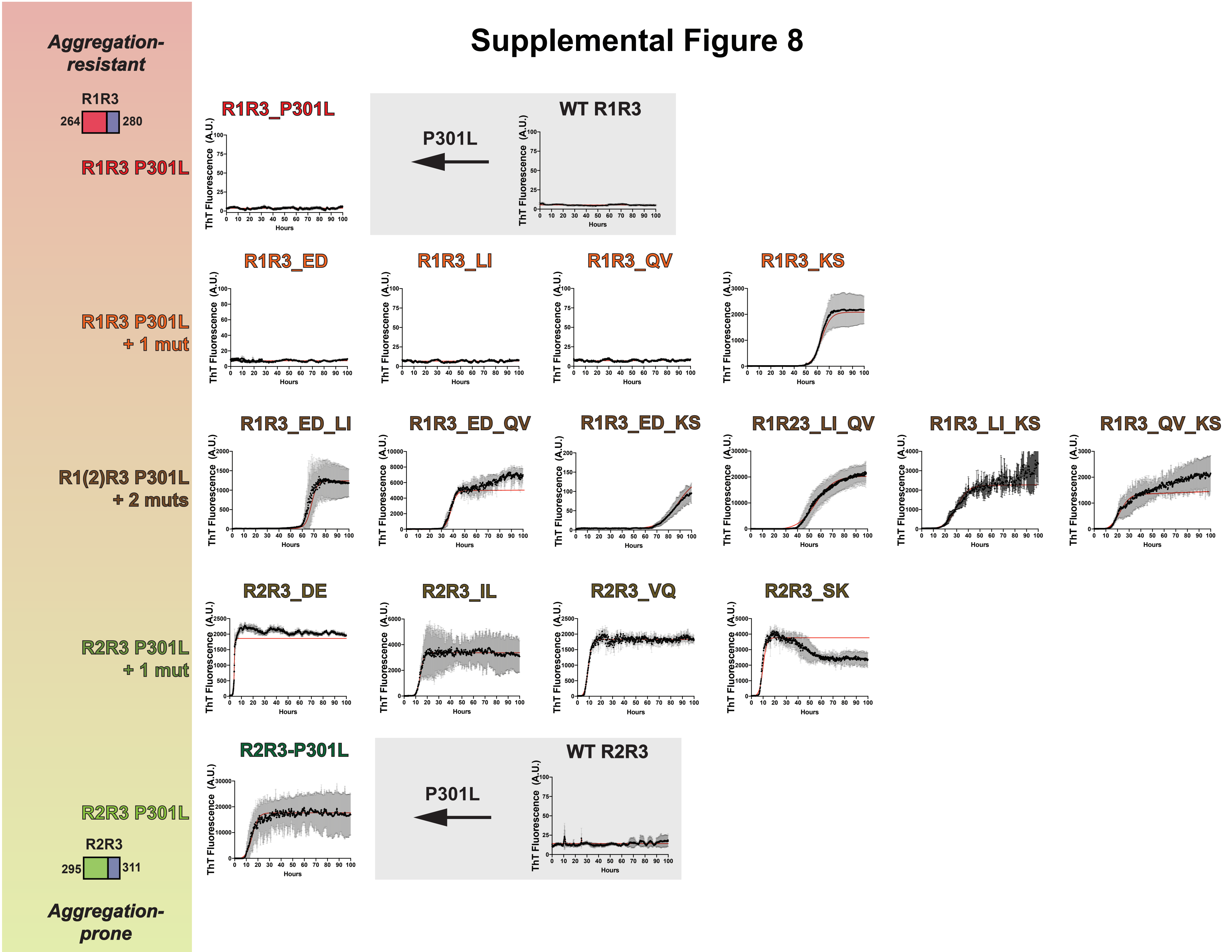

### Supplemental Figure 9

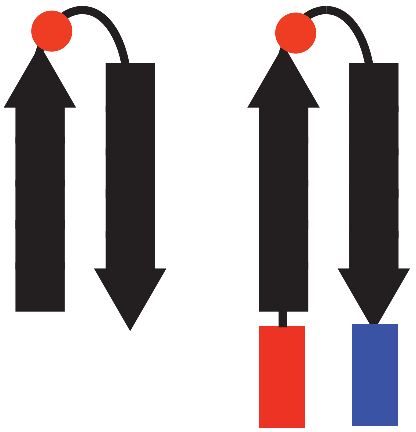

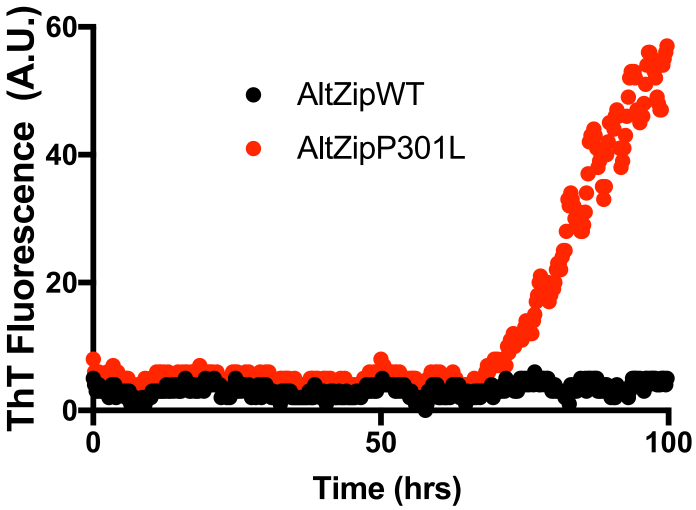

### Supplemental Figure 10

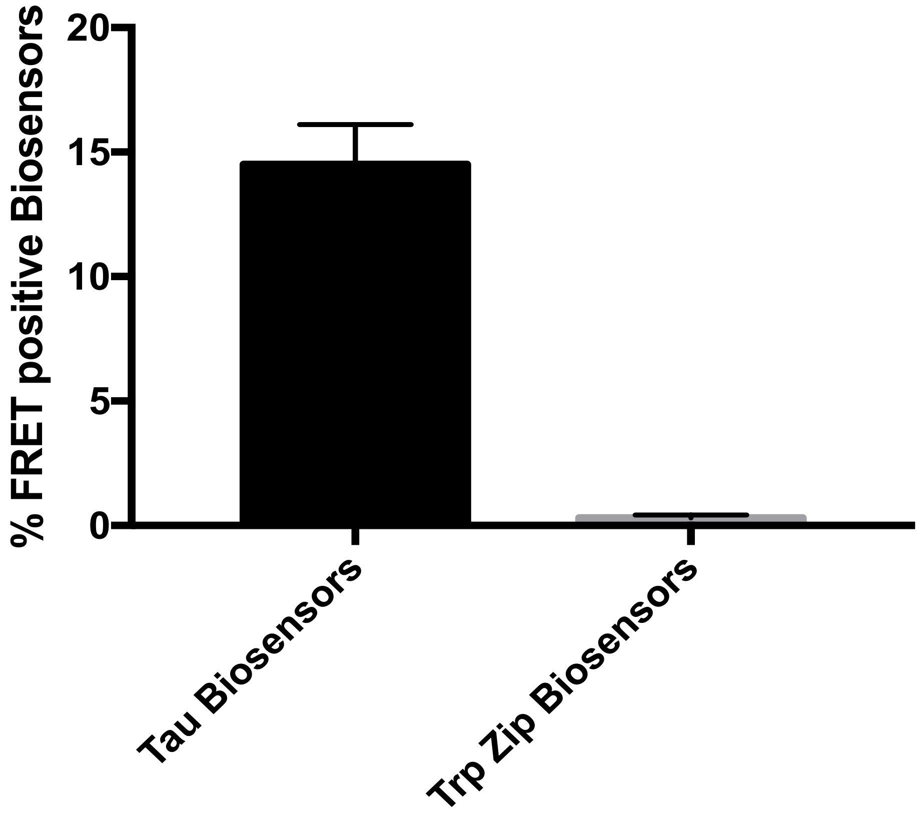
