## Supplemental Figure Legends for "Tau local structure shields amyloid motif and controls aggregation propensity"

**Supplemental Figure 1. Intramolecular FRET measurement for $M_i$ and $M_s$ tau.** (**a**) Cartoon schematic of tau used for intramolecular FRET studies colored according to repeat domain (repeat 1 = red; repeat 2 = green; repeat 3 = blue; repeat 4 = purple) and highlighting the location of the cysteines used for attachment of donor and acceptor dye that flank the $^{305}$VQIVYK$^{311}$ motif. (**b**) Histogram of FRET efficiency measurements for $M_i$ and $M_s$ at 25ºC and following heating at 95ºC. Each measurement was performed in replicates.

**Supplemental Figure 2. TEM of tau and tau RD aggregates.** Transmission electron microscopy images of each protein sample from previous ThT experiments after 5 days (**Figure 1 c-e**).

**Supplemental Figure 3. Comparison of consensus crosslinks between WT, P301L and P301S tau RD.** (**a**) SDS-PAGE/coomassie of control and DSS reacted samples, indicating the protein remains a monomer following cross-linking (indicative of intra- and not inter-crosslinks) for the WT and mutants at each temperature. (**b**) Total consensus crosslinks (crosslinks found in all 5 of 5 replicates) in WT (black) and P301L (dark gray) tau RD at 37ºC (yellow), 50ºC (orange) and 75ºC (red). (**c**) Venn diagrams showing overlap between WT (purple) and P301L (orange) tau RD consensus crosslinks acquired at 37ºC, 50 ºC and 75ºC. (**d**) P301S tau RD samples were incubated at 37ºC, 50ºC or 75ºC for one hour. After cross-linking, trypsin fragmentation, and LC-MS/MS analysis, consensus cross-link patterns (circles) are shown as contact maps color coded by average frequency across replicates. The theoretical contacts are shown in the background as gray circles. Short range crosslinks within the N-term (blue), C-term (red) and contacts across N- to C-term (purple) are shown as sectors. The x- and y-axis are colored according to repeat number as in Figure 1. The dashed boxes define inter-repeat crosslinks observed between repeat 2 and repeat 3.

**Supplemental Figure 4. Quantification of false discovery rate in crosslink identification.** Representative true positive (red) and false positive (green) distribution plots separated by score-id to calculate false discovery rates (FDRs) for each XL-MS dataset. A cutoff with score=25 (vertical dashed lines) is applied to estimate FDR (**Methods**). FDR is a qualitative measure calculated as ([Number of false positives with score ≥ 25] / [total number of crosslinks with score ≥ 25]) or (1 / [total number of crosslinks with score ≥ 25]) if there is no false positive above score 25.

**Supplemental Figure 5. ROSETTA-based simulations of tau RD reveal possible hairpin structures.** The structural ensembles are shown as a distribution of total energy of each model and radius of gyration for (**a**) tau RD CS-ROSETTA and (**b**) tau RD *ab initio*. (**c**) Secondary structure models of the repeat2-repeat3 interface ($^{295}$DNIKHVPGGGSVQIVYK$^{311}$) predicted from CS-ROSETTA (**d**) or *ab initio* shown in cartoon and colored by repeat domain. The amyloid motif $^{306}$VQIVYK$^{311}$ is in blue, the leading repeat-2 sequence is green. Alignment of the structural ensemble to the template model highlights the broad diversity of structures produced by the simulation. The data are shown as Rosetta score versus $c\alpha$-$c\alpha$ root mean square deviation (rmsd) plots. (**e**)

Pairwise alignment of template hairpin for R1R2, R2R3, R3R4 and R4R’ region to each member in the ensemble is shown as a function of Rosetta score revealing all hairpin containing models. A 5 Å$^2$ rmsd was used as a threshold to define similarity to a template hairpin. The regions and sequences used in the alignment are shown on the left and colored according to the repeat domain as in Figure 1. (**f**) Positions of disease-associated mutations (red spheres) mapped onto the $^{306}$VQIVYK$^{311}$-containing *ab initio* $\beta$-hairpin structure.

**Supplemental Figure 6. Energetics and hairpin content in ROSETTA structural ensembles.** (**a-d**) Ensemble-wide distance distribution of end-to-end distances between residues 264-280, 295-311, 327-347 and 359-375 across all inter-repeat hairpins from *ab initio* and CS-ROSETTA models. A 15 Å c$\alpha$-c$\alpha$ distance was used as a threshold to define a collapsed hairpin-like structure.

**Supplemental Figure 7. An amyloidogenic hexapeptide motif is sufficient to aggregate spontaneously.** The aggregation of hexapeptides representing recognized amyloidogenic regions of tau ($^{275}$VQIINK$^{280}$ and $^{306}$VQIVYK$^{311}$) or the N-terminal flanking sequences ($^{264}$ENLKHQ$^{269}$ and $^{295}$DNIKHV$^{300}$) were studied using ThT fluorescence. The signals are shown as the average of triplicates with standard deviation. The $^{264}$ENLKHQ$^{269}$ and $^{295}$DNIKHV$^{300}$ peptides yielded no detectible ThT signal change (less than 2-fold ratio to background signal) over the course of the experiment (see Supplemental Table 1). ThT signals are shown as average of triplicates with standard deviation and are normalized to the maximum for each condition.

**Supplemental Figure 8. Aggregation kinetics of chimeric peptides derived from R1R3 and R2R3 sequences.** The ThT curves for 16 peptides containing single and double mutations are shown as an average from technical triplicates with standard deviations. The mutants are colored according to repeat composition R1R3-P301L (red), R1R3-P301L single mutants (+1 mut, orange), R1(2)R3-P301L double mutants (+2 muts, brown), R2R3-P301L single mutants (+1 muts, dark green) and R2R3-P301L (green). The R1R3-WT and R2R3-WT peptide controls are shown in grey. The data were fit to a non-linear regression model (red line) to derive $t_{1/2}$ values shown in Figure 6.

**Supplemental Figure 9. A salt bridge stabilizes and rescues P301L mutant spontaneous aggregation.** An alternative method to stabilize the ends of predicted β-hairpin was used in addition to a tryptophan zipper. Two additional glutamic acid and lysine were placed at the flanking N- and C-termini respectively (**Table 1**). Doing so delayed aggregation of R2R3-P301L ($t_{1/2}$ = 7 hours; **Figure 4**) by over an order of magnitude ($t_{1/2}$ > 70 hours).

**Supplemental Figure 10. Tau biosensor cells with enhanced local structure are resistant to being seeded by exogenous aggregates.** Tau biosensor cells were constructed with a tryptophan zipper as what was used in the trp-R2R3-P301L-trp peptide fragment, but in the context of tau RD-CFP/YFP (**Methods**). When transduced with exogenous tau aggregate material, the TrpZip encoded biosensors showed a significant reduction in capacity for seeding. % FRET positive cells were measured using Flow Cytometry in triplicates with at least 10,000 cells per condition.

**Supplemental Table 1. Summary of ThT experimental values for tau, tau RD and peptide aggregation experiments.**

**Supplemental Table 2. Summary of calculated FRET efficiencies for free Alexa-488 and Alexa-647 dyes at 37ºC, 50ºC and 75ºC.**

**Supplemental Table 3. Summary of technical replicate XL-MS data for WT, P301L and P301S tau RD at 37ºC, 50ºC and 75ºC.**

**Supplemental Table 4. Summary of consensus crosslink pairs for WT, P301L and P301S tau RD at 37ºC, 50ºC and 75ºC.**
